# Electrical Stimulation of Human Mesenchymal Stem Cells on Conductive Chitosan-Polyaniline Substrates Promotes Neural Priming

**DOI:** 10.1101/2022.11.14.516447

**Authors:** Behnaz Sadat Eftekhari, Dawei Song, Paul A. Janmey

## Abstract

Electrical stimulation (ES) within conductive polymer substrates has been suggested to promote the differentiation of stem cells toward a neuronal phenotype. The use of conductive scaffolds in tissue regeneration provides a unique and attractive new option to control the amount and location of ES delivery. Scaffold stiffness has also been shown to be an important regulator of stem cells’ behavior and fate. Therefore, to improve stem cell-based regenerative therapies, it is essential to characterize the simultaneous effects of electroconductive substrate stiffness and electric field stimuli on stem cell fate processes. In this study, biodegradable electroconductive substrates based on chitosan-polyaniline (CS-g-PANI) were fabricated with different stiffnesses. Human mesenchymal stem cells (hMSCs) seeded on these scaffolds were electrically stimulated for 14 days with 100 mV/ cm (20 min every day). For hMSCs cultured on soft conductive scaffolds, a morphological change with significant filopodial elongation was observed after 2 weeks of electrically stimulated culture. Compared with stiff conductive CS-g-PANI scaffolds and non-conductive CS scaffolds, for soft conductive CS-g-PANI scaffolds microtubule-associated protein 2 (MAP2) and neurofilament (NF-H) expression increased after application of ES. At the same time, there was a decrease in the expression of the glial markers glial fibrillary acidic protein (GFAP) and vimentin after ES. Furthermore, the elevation of intracellular calcium [Ca^2+^] during spontaneous, cell-generated Ca^2+^ transients further suggested that electric field stimulation of hMSCs cultured on conductive CS-g-PANI substrates can promote a neural-like phenotype. Our findings propose that the combination of the soft conductive CS-g-PANI substrate and ES is a promising new tool for enhancing nerve tissue engineering outcomes.

## 1. Introduction

Nerve injuries leading to reduced motor or sensory function affect millions of people in the world each year, and because of the complexity of the nervous system and its limited regenerative capacity, achievement of efficient therapeutic strategies has been difficult [1, 2]. The rapid development of regenerative medicine has demonstrated that engineered nerve scaffolds can overcome the limitations of previous therapies and promote neural recovery. However, it remains highly challenging to design and fabricate an ideal scaffold with suitable properties that match those of native tissues [3].

The existence of endogenous electric fields (EF) has been identified in several tissues, such as neural, cardiac, and skeletal muscle tissues [4]. Several recent studies have revealed the important role of electrical stimulation (ES) in neurite extension, as well as the promotion of neurotrophin secretion and myelin gene expression by stem cells, which lead to neural differentiation and proliferation [5, 6]. However, the exact mechanism of how ES affects biological systems remains unclear. Nevertheless, it has been shown that ES pathways can regulate various cellular events including membrane enzyme protein function (Na+/K+ATPase and Ca^2+^ ATPases), alteration of cell-surface receptor formation, cytoskeleton reorganization, changes in intracellular calcium ion concentration, transporting ion channel activation, and cell shape change [7-9].

Considering the biological importance of ES, electroconductive scaffolds have been developed to control the localization, amount, and duration of ES delivery to cells as well as to support tissue growth and regeneration [10]. Among various classes of conductive materials, polyaniline (PANI) is one of the most promising biomaterials owing to its excellent conductivity, high environmental stability, suitable mechanical properties, and flexibility in processing [11]. However, poor solubility and degradation properties of PANI restrict its application in its pristine form [12]. In contrast, natural polymers have been utilized as tissue engineering scaffolds because of their proper biocompatibility and controllable biodegradability; however, they are generally non-conductive [13]. Several studies reported promising neural regeneration using chitosan based-scaffolds [14-17]. Jiang et al. have successfully employed a chitin/CM-chitosan nerve graft to repair a 10-mm defect in adult male Sprague Dawley rat [18]. Chitosan (CS), a derivative of the natural polymer chitin, is recognized as a promising biomaterial for tissue engineering scaffolds due to its biodegradability, antimicrobial activity, and low toxicity and immunogenicity [19]. Therefore, the synthesis of a composite using PANI and CS has potential to produce an advantageous water-soluble conductive material with good electroconductivity.

On the other hand, the use of human mesenchymal stem cells (hMSCs) has been extensively investigated for neural differentiation applications, because of their ease and low cost for isolation, immune compatibility, abundance in tissues, and ability to release neurotrophic factors [20, 21]. However, it is still challenging to find a well-established matrix that provides a biomimetic environment for hMSCs to efficiently direct their differentiation into a neuronal lineage. It has been reported that an external ES applied to hMSCs changed their alignment and morphology [22], improved cell migration [23], increased cell proliferation [24], and upregulated neural gene expression [25]. Park et al. demonstrated that electrically stimulated hMSCs co□cultured with mature neuronal cells produced the highest level of neurite outgrowth (>150□mm) and smaller cell body sizes [26]. In particular, electrical stimulation can influence the activation of Voltage-Gated Calcium Channels (VGCC) leading to an increase in intracellular Ca2+ influx. [27]. Specifically for bone marrow-derived hMSC stimulation, increased levels of neuronal marker gene expression are associated with CREB-mediated phosphorylation [28]. In view of the above, we hypothesized that ES stimulation within a conductive substrate would improve the neural priming of stem cells that we observed with hMSCs in CS-g-PANI hydrogels in the absence of an external EF. In a previous study we showed that the conductive CS-g-PANI blend hydrogel can promote the neurogenic phenotype of MSCs, likely due to electrical signals endogenously generated by the cells [29].

Moreover, substrate elasticity has been reported to regulate phenotype commitment of human mesenchymal stem cell (hMSCs) since the pioneering work of *Engler et al*. [30]. It has been shown that culture substrate stiffness can change the assembly of F-actin fibers within the cell and alter cell morphology and spreading [31]. Also, because the linker of the nucleoskeleton and cytoskeleton (LINC) complex bridges the cytoskeleton and nucleus to transmit mechanical forces to chromatin [32], the substrate stiffness might modulate genomic structure, expression of downstream genes, and eventual cell fate and behavior [33]. Therefore, it is essential to examine the effects of both ES and hydrogel elasticity on neural differentiation of hMSCs. However, there is a concern that scaffold with different stiffnesses might exhibit different electrical conductivities, making it difficult to interpret their isolated effects on stem cell fate.

To address the question raised above, we developed a method to control hydrogel stiffness, independent of electrical conductivity. In the present study, we synthesized a conductive composite by grafting PANI onto a CS network and then develop CS-g-PANI hydrogels with different stiffnesses and the same range of conductivity. The hMSCs were seeded on CS-g-PANI and CS substrates (non-conductive substrates serving as control samples) and were stimulated by an electric field (100 mV/cm) for 20 min per day for 14 days in culture. The cells were also cultured on both substrates in the absence of ES in order to determine the neurogenic potential of electrical stimulation and substrate stiffness.

## 2. Experimental

### 2.1 Synthesis of chitosan-grafted-polyaniline (CS-g-PANI)

The CS-g-PANI copolymer was synthesized by radical polymerization following the procedure reported by Marcasuzaa et al [34]. CS (DD=80, medium molecular weight, Sigma-Aldrich, St. Louis, MO, USA) solution was prepared by mixing 0.1g of CS in 10 ml 0.1 M acetic acid (Merck, Darmstadt, Germany). The solution was stirred for 12 h at room temperature to obtain a clear solution. 5.0 mL of aniline (St. Louis, MO, USA) solution was then mixed in 20 ml 1 M aqueous hydrochloric acid (HCL) (Merck, Darmstadt, Germany) solution at room temperature. 10 mL of the CS solution and 25 mL of the aniline solution were mixed on a magnetic stirrer for 30 minutes at 0-5 °C. A solution of 0.1 M ammonium persulfate (APS) (St. Louis, MO, USA) prepared in 1 M HCl solution was added dropwise into the CS-g-PANI solution in an ice bath. The stirring was continued overnight under N2 atmosphere and a crude product (dark green in color) was obtained. Neutralization of the reaction mixture was performed using 5 M NaOH, and the obtained polymer was precipitated into excess ethanol. The resultant precipitate was filtered and washed with N-methyl-2-pyrrolidone (NMP) (Sigma-Aldrich, St. Louis, MO, USA) to remove any PANI ungrafted onto CS. After washing with acetone and drying in vacuum at 50 °C the desired CS-g-PANI copolymer was produced.

### 2.2 Preparation of CS and CS -g-PANI hydrogel substrates

To fabricate a composite hydrogel by a gel-casting technique, the required amounts of CS solution (in 0.1 M acetic acid and glycerol (Sigma-Aldrich, St. Louis, MO, USA) in a ratio of 3:2) and CS-g-PANI composite dispersion (in water) were mixed together with a mechanical stirrer at 1000 rpm for 1 h. Pure CS solution was also prepared by dissolving 0.1 g of chitosan in 10 mL of a mixture of 0.1 M acetic acid and glycerol ((Sigma-Aldrich, St. Louis, MO, USA) in a ratio of 3:2).

For the preparation of stiff hydrogel films, both composite solution and CS solution were allowed to dry at 60 °C in a hot-air oven, the films were peeled off and neutralized with 5 N NaOH. Then, these films were washed thoroughly and kept in a vacuum oven for further drying. The corresponding soft hydrogel samples were obtained after pouring the CS-g-PANI composite or CS solutions in Petri dishes (with a diameter of 3.5 cm) and neutralization with 5 N NaOH. The samples were thoroughly washed with deionized water until neutrality and complete removal of sodium hydroxide and salts; then, the samples were stored in deionized water.

### 2.3 Characterization of CS-g-PANI and CS substrates

#### 2.3.1 Fourier-transform infrared spectroscopy (FTIR) spectra

The FTIR spectra of CS-g-PANI and CS hydrogels were recorded with a Thermo Nicolet Nexus FTIR spectrometer in the transmittance mode in the frequency range of 4000– 400 cm^-1^.

#### 2.3.2 Atomic force microscopy (AFM measurement)

AFM tapping mode was performed with a NanoWizard® 4 BioScience AFM (JPK BioAFM, Bruker Nano GmbH, Germany) to evaluate the surface of our hydrogels. A non-conductive silicon nitride cantilever (DNP-10, BRUKER, USA) with a spring constant of 0.35 N/m was used. The surface of the sample (up to 20 μm^2^) was scanned at a scan rate of 0.05 Hz and setpoint force 0.5 nN. The standard software of the instrument (JPKSPM Data Processing) was utilized for image analysis. Before roughness values (R_a_) measurement, plane fit and line leveling filters were applied to the AFM images; thereafter, R_a_ was calculated based on 3 images per sample and reported as mean ± SD.

#### 2.3.3 Electrical conductivity

The electrical conductivity of each prepared sample was measured by applying the two-point probe method (Keithley, model 7517A). The electrical conductivity (σ) was measured by passing a constant current through the outer probes and recording the voltage via the inner probes. The conductivity of the samples was calculated as follows:

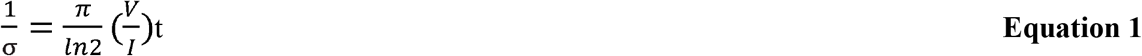

where *σ*, I, V, and t indicate, respectively, the electrical conductivity, applied current, voltage, and the sample thickness (200 μm).

#### 2.3.4 Swelling ratio

Three samples of each substrate were prepared with the same size and shape and were then weighed (W_0_). Next, they were incubated in PBS solution at 37 °C. After 24 h, the excess liquid was removed by filter paper and the swollen sample was again weighed (W_1_). W_0_ and W_1_ were each measured three times. The swelling ratio was calculated by the following equation:

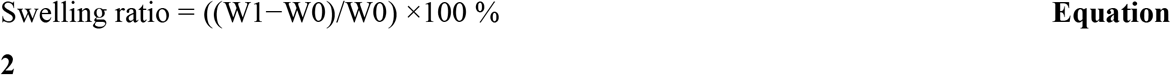

#### 2.3.5 Biodegradation assay

To evaluate the biodegradation rate of the scaffolds, CS and CS-g-PANI samples with the same initial weight (W_0_) were incubated with 1 mg mL^−1^ collagenase type I in PBS (pH= 7.4) and placed on an orbital shaker at 100 rpm and 37 °C. At predetermined time intervals, a sample was taken out, and its dried weight was measured (W_1_). The biodegradation stability of prepared scaffolds was assessed by calculating the percentage of the remaining weight after enzymatic degradation, via

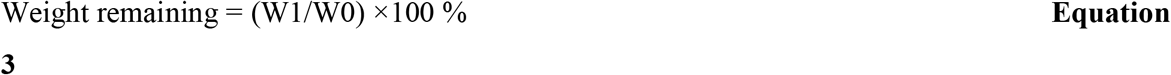

#### 2.3.6 Mechanical testing

A rotational KINEXUS rheometer (Malvern Instruments) with parallel plates was used to measure the mechanical properties of the scaffolds. Both CS and CS-g-PANI samples were cut using a steel punch into cylindrical disks of 8 mm diameter. A plate of 8 mm diameter and a gap of 60-150 µm was used for the parallel-plate rheometer. Sandpapers were attached to the plates to avoid sliding between the samples and plates. A Peltier device incorporated into the bottom plate was used to control the sample temperature at 25 °C.

A series of mechanical tests were conducted to define the mechanical properties of the samples. First, to compare the stiffness of different samples in the resting state, the shear storage modulus was measured by applying a 2% oscillatory shear strain at a frequency of 1 Hz for 3 minutes. Second, to test the frequency dependence of the sample properties, shear moduli at 2% oscillatory shear strain were measured at frequencies between 0.01 and 10 Hz. Third, the samples’ shear moduli were measured at shear strains between 0.5 and 50% at 1 Hz. Fourth, stress relaxation of the samples in response to constant shear deformation was measured by applying a 5% steady shear strain and recording the resulting shear stress as a function of time. Fifth, stress relaxation after compressing the samples by 30% was measured from the resulting normal force as a function of time. We note that in shear relaxation tests, a sample preserves its volume, and stress relaxation is an indication of the viscoelastic response of the sample. In contrast, in compression relaxation tests a sample can change its volume, because solvent can be squeezed out of the compressed sample. Thus, stress relaxation is mainly a consequence of the poroelastic and viscoelastic responses of the sample.

Finally, to test if application of an electric field altered the mechanical properties of the conductive polymer, a customized torsion pendulum based on a previous design [35] was constructed using non-conductive building blocks and glass slides as the sample holders. The chitosan-PANI substrate was placed between the two circular glass plates of the torsion pendulum, and two copper wires were placed in contact with free edges of the substrate at opposite sides of the disk-shaped sample. The shear storage and loss modulus were calculated from the frequency and damping rate of the free oscillations induced by a small impulse to the pendulum arm provided by a puff of air [36]. Measurements were made before and immediately after application of 18V across the 8 mm wide sample.

### 2.4 Cell- substrate interaction

#### 2.4.1 Evaluation of biocompatibility

Before being seeded with cells, the samples were repeatedly treated with 75% ethyl alcohol and UV light for sterilization. To apply ES on cells, a home-built electric field device was used as illustrated in Figure S1. Briefly, two electrode wires with 1 cm spacing were placed on the two ends of sterile tissue culture plates. The round sample was positioned at the center of the prepared chamber. Copper wires were used to connect the platinum electrodes and electric generator (FLIR Systems Extech 382202 Digital Single Output DC Power Supply, USA). Sterilization of the assembled chamber was performed before use by placing under UV light for 1 h. hMSCs (1 × 10^5^/ml) were seeded onto CS and CS-g-PANI scaffolds and cultured at 37 °C for 24 h to allow cell attachment. A constant electrical field of 100 mV cm^−1^ was applied across the two electrodes for 20 min (at 37 °C with 5% CO_2_) daily for 2 weeks to stimulate the cells.

#### 2.4.2 Cell viability and proliferation of hMSCs on hydrogels

To optimize the biocompatible ES intensity with hMSCs, different electric field intensities (50, 100, 150 and 200 mV/cm) were applied to the cells seeded on scaffolds from the second day of cell seeding (for simplicity, the day on which the ES was first applied was taken as the first day). In order to stain the live and dead cells, calcein-AM and ethidium homodimer (Thermofisher, USA), were used, respectively. Following the electrical stimulation at each time point, the culture media was taken out of the samples. The cells were incubated with 1 mL of calcein-AM (25 μg/mL) and 1 mL of ethidium homodimer (10 μg/mL) for 30 min at room temperature. Subsequently, the samples were washed thrice with PBS and observed under a fluorescence microscope (Nikon, LV 100D, Japan). After optimization of ES intensity, cell viability on the CS and CS-g-PANI hydrogel films was determined via the MTT (Thermofisher, USA) assay at 3, 7 and 14 days with/without optimized EF. After culturing for respective time points, cells on the substrates were washed three times with PBS, followed by the addition of 500 μL of MTT (0.5 mg/mL) reagent prepared in DMEM (Thermofisher, USA) for 4 h at 37 °C. Then 300 μL of DMSO was added and incubated for 10 min to dissolve the formed formazan precipitates. Finally, the optical density (OD) was measured using a spectrophotometer at a wavelength of 570 nm. The absorbance value was determined using the following formula:

Absorbance value (OD) = Average OD of the samples – Average OD of the negative control, where the cell culture media without substrates and cells was considered as a negative control. Cells cultured on the plastic surface of the cell culture plate served as positive control (considered as 100% cell viability). All data of the test were expressed as means ± standard deviation (SD) for n = 4. The Kolmogorov–Smirnov test was applied to investigate the normal distribution of each group of samples. Finally, a one-way ANOVA and Duncan test (GraphPad Prism 7 software) was performed to compare the data (P<0.05 was considered a significant difference). In addition, nuclei staining was performed to identify cell proliferation on the substrates with and without ES. Following cell culture, the cells on the CS and CS-g-PANI hydrogel films were fixed with paraformaldehyde for 15 min and incubated with 1 mL of DAPI (20 μg/mL in PBS) for 10 min, and then imaged. The fluorescence images of the cells cultured in different conditions were used to compare the difference in cell number by using the Image J software.

#### 2.4.3 Morphology of hMSCs on hydrogels

To evaluate the effect of ES and substrate stiffness on cell morphology and attachment, F-actin staining was carried out. The hMSCs were cultured on different substrates at 5000 cells/cm^2^. After 72 hours, cells were rinsed with 1x PBS, fixed with 4% polyformaldehyde (PFA) (Sigma-Aldrich, St. Louis, MO, USA) and permeabilized with 0.1% Triton X-100. After rinsing with PBS three times, Alexa-Fluor 647 phalloidin (Invitrogen, USA) and DAPI (Sigma-Aldrich, St. Louis, MO, USA) were added to samples for 1 hour in the dark. Cells were observed and photographed under an inverted fluorescence microscope (Eclipse Ti-S, Nikon).

#### 2.4.4 Neural differentiation of MSCs on hydrogels

hMSCs were seeded on a six-well culture plate at a density of 5 ×10^5^ cell/ml with 1 ml of neural induction medium containing serum-free DMEM/F12 supplemented with 200 μM ascorbate and 2 mM glutamine (Invitrogen, USA) along with 2% B-27 (Invitrogen, USA) for 14 days. The induction medium was replaced every 2 days. The differentiation process lasted for 14 days, and then the cells were analyzed by immunofluorescent staining.

#### 2.4.5 Immunofluorescence staining

After 14 days from neural induction with ES, the cells seeded on substrates were rinsed with PBS and fixed with 4% PFA in PBS (pH 7.4) for 20 min. In the next step, cells were permeabilized in a solution containing 0.5% Triton X-100 in PBS for 10 min. To block non-specific antibodies, cells were incubated with 10% normal goat serum for 1 h at room temperature (0.05% Tween 20 and 1% (w/v) bovine serum albumin (BSA)). Detection and characterization of neuronal cells were carried out by staining the cells with antibodies against GFAP, MAP2, NFL and vimentin. Antibodies to GFAP (1:100; Abcam), MAP 2 (1:50; Millipore), NFL (1:100 dilution), and vimentin (1:100 dilution), were added to the cell medium for staining overnight at 4°C. After cells were washed with PBS, the secondary antibody, including Alexa 488-conjugated goat anti-mouse and anti-rabbit antibodies, was added to cells and incubated for 1h. Finally, the nuclei of cells were counterstained with DAPI. Fluorescence images were taken by an Olympus BX51 fluorescence microscope. To determine the percentage of positive cells, the occurrence of neuronal markers (NFL or MAP-2 positive cells) and glial markers (GFAP positive cells) was counted and calculated as a percentage of total DAPI positive nuclei in 5 random visual fields.

#### 2.4.6 Neurite measurement

To evaluate the influence of the conductive CS-g-PANI substrate, electrical stimulation, and hydrogel stiffness on hMSCs, neurite length and the percentage of differentiated cells were measured using ImageJ software. Neurite length was defined as the straight-line distance from the tip of the neurite to the cell body. The percentage of differentiated cells was counted within each microscope field. For each photo, 3−5 views were selected, and at least three photos of each sample were analyzed.

#### 2.4.7 Intracellular calcium imaging

To determine changes in Ca^++^ concentration in differentiated hMSCs, fluo-4-acetoxymethyl ester (Fluo-4 AM, 5 Mm, Invitrogen, USA) was added to the medium after day 14 of neural induction with electrical stimulation. Briefly, medium was aspirated and washed with Hank’s Balanced Salt Solution (HBSS, Sigma, USA) and Tyrode’s solution. Cells were treated with staining solution containing Fluo-4, Pluronic acid, and Tyrode’s solution at room temperature for 45 minutes. After discarding staining solution, cells were washed with Tyrode’s solution and HBSS, and incubated at 37°C for 15 minutes in induction medium. Time-lapse imaging of the calcium level in live hMSCs was done using Nikon LV100D fluorescence microscopy. The recordings were initiated at the start of the ES exposure and then continued for 15-20 min. During microscopic imaging, the cells were excited at 495 nm and emission captured at 515 nm. The change in the fluorescence intensity was normalized with the baseline fluorescence value for individual cells (region of interest).

### 2.5 Imaging and Statistical analysis

Morphometric and quantitative analyses of images were performed in Image J Pro Plus or Origin Pro. Statistical analyses were performed using SPSS. The data are presented as the means ± SD, and p < 0.05 was considered statistically significant. Statistical significance was determined by one-way analysis of variance (ANOVA) followed by Tukey’s multiple comparison tests.

## 3. Results and discussion

Previous studies show that the biophysical characteristics of scaffolds, such as topographic features at the micro- and nanoscale [37], mechanical properties [38], and conductivity [39], can regulate the behavior of the mammalian cells [37]. Jin et al. reported the capability of electrical stimulation to recapitulate the brain-like environment, which can in turn facilitate neuronal differentiation of the stem cell [40]. On the other hand, previous studies have shown that soft substrates with elasticities commensurable to the elasticity of the brain tissue can enhance stem cell differentiation into neural like cells [41, 42]. Such microenvironment cues play a critical role in cellular fate. In order to create a biophysically simulated microenvironment for the cellular differentiation, the present study utilized a conductive biodegradable scaffold with two different stiffnesses to apply electric field (100 mV/ cm) stimuli to hMSCs, and to examine the synergistic effect of substrate stiffness and electrical stimulation. The analysis of cell viability, morphology, and protein expression allowed us to identify the cell fate under different proliferative/differentiative conditions. Furthermore, the cell functionality and differentiation status of the electrically stimulated cells was evaluated.

### 3.1 FTIR spectrum

In this study, the conductive CS-g-PANI composite hydrogels with different stiffnesses were prepared by a two-step process. In the first step, a conductive composite was synthesized by grafting PANI to CS network through oxidative-radical polymerization. In the second step, CS-g-PANI hydrogels was formed by a gel-casting method. The non-conductive CS hydrogels with different stiffnesses were also made by the same method to be used as the control samples (Figure 1A). FT-IR spectroscopy was employed to obtain information about the structure and different types of interaction in the hydrogel matrix. The FTIR spectra of CS and CS-g-PANI substrates are demonstrated in Figure 1B. The CS samples spectrum presented an expected absorption peak at 3335 cm^-1^ due to –NH2 stretching and the bands 1644–1380 cm^-1^ due to – NH2 bending. The band located at 2882 cm^-1^ is assigned to the C–H stretching mode in chitosan. The peak at 1260 cm^-1^ corresponds to asymmetric stretching of the C–O–C bridge, while the peak at 1148 cm^-1^ shows the vibration of the C–O stretching [43]. These peaks are characteristics of its saccharide structure. The CS-g-PANI infrared spectrums revealed all the characteristic peaks corresponding to chitosan and PANI. In this spectrum, the peak observed at 3340 cm^-1^ exhibits –NH2 stretching with amino groups in chitosan. The bands observed at 1637 cm^-1^ could be from the stretching mode of C=O, and are generally due to saccharides. The appearance of C=C stretching modes of the quinoid ring and benzenoid ring at 1590 cm^-1^ and 1391 cm^-1^, respectively, is a characteristic of PANI [44, 45].

**Figure 1:**
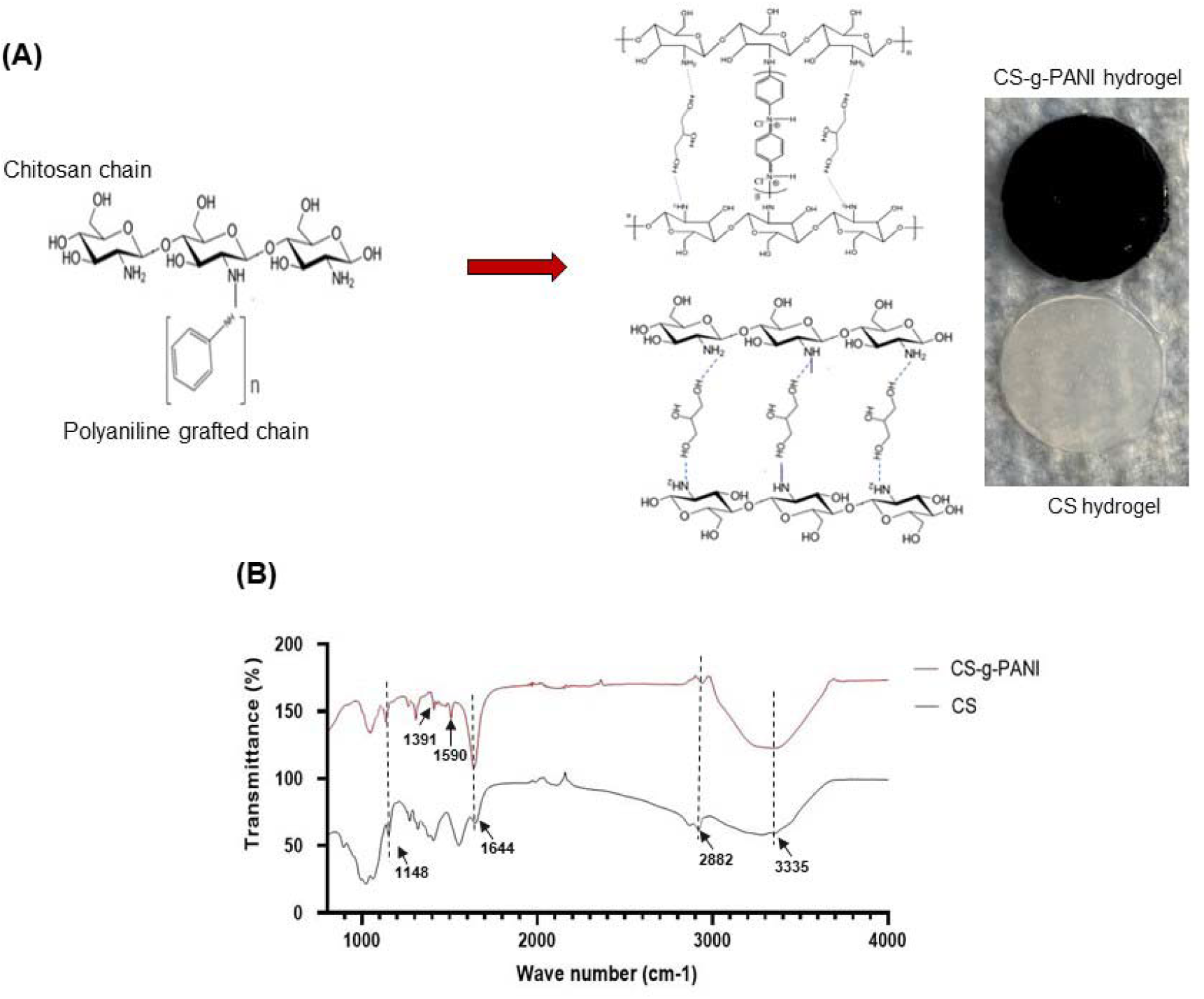
(A) Schematic representation of synthesis of chitosan-grafted-polyaniline (CS-g-PANI) copolymer and the formation of the CS and CS−PANI hydrogel nanocomposite film. (B) FTIR spectrum of CS and CS-g-PANI.

### 3.2 Atomic force microscopy (AFM) imaging

AFM height images and their 3D representations for non-conductive CS and conductive CS-g-PANI samples are shown in Figure 2A-H. According to the topography images, CS-g-PANI composite hydrogels had a relatively rougher surface topology than CS hydrogels, presumably due to the grafting of PANI into the CS network after polymerization. The height images of the soft hydrogels (Figure 2G-H) showed that these samples had higher surface roughness as shown by the respective representative surface profiles. For these soft samples, the mean roughness values were R_a_ = 180 ± 6 nm for CS hydrogels and R_a_ = 314 ± 4 nm for CS-PANI. In contrast, for stiff samples (Figures 2E-F) the mean roughness values were R_a_ = 29 ± 7 nm for CS hydrogels and R_a_ = 38 ± 2 nm for CS-PANI.

**Figure 2:**
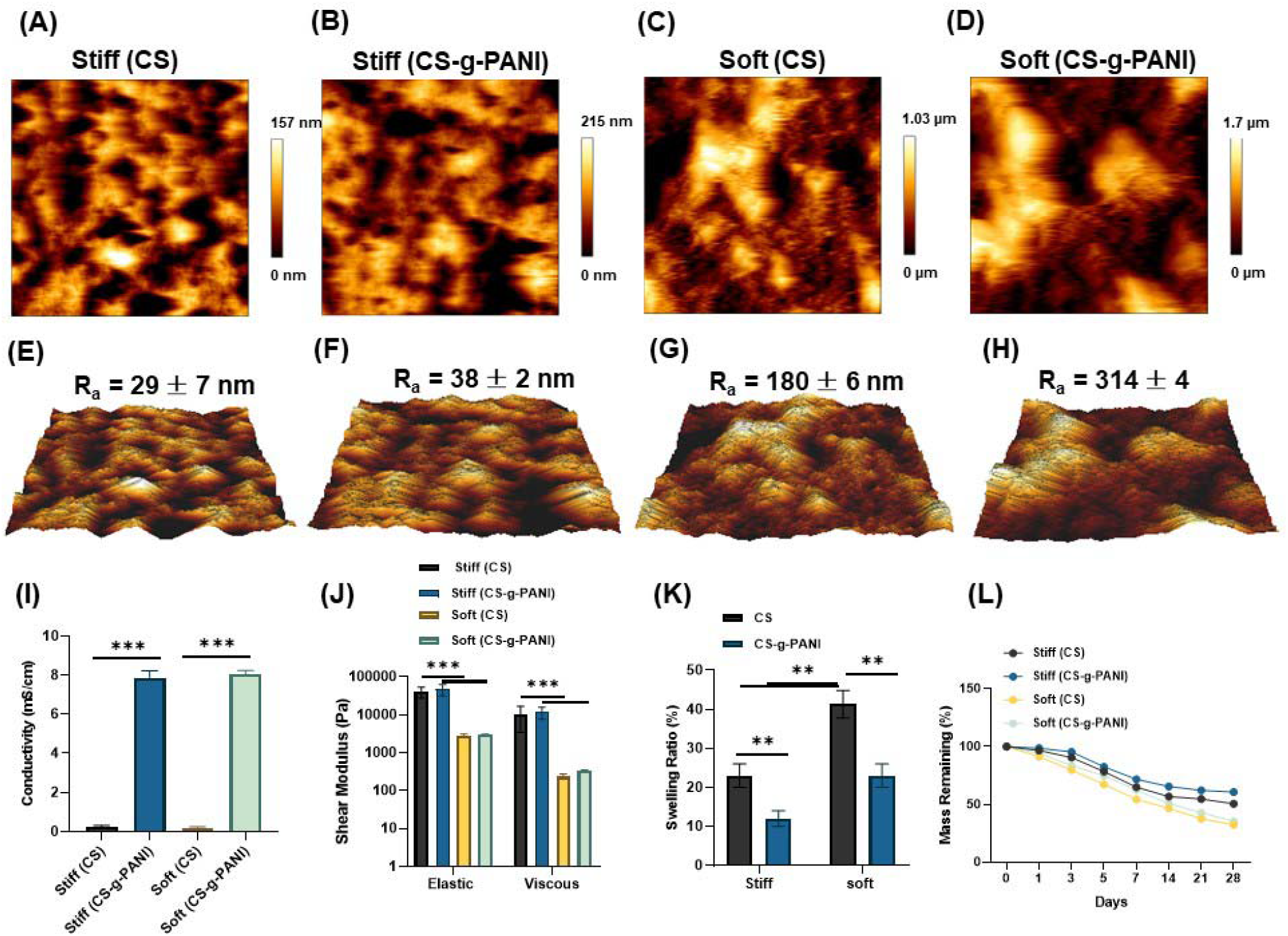
(A–D) AFM height images of the stiff and soft CS and CS-g-PANI samples. (E–H) 3D representation and relative height roughness (Ra). (I) Electrical conductivity of pure CS increased after grafting PANI for both samples. (J) Swelling ratio of the CS and CS-g-PANI hydrogels. (K) The elastic and viscous shear moduli of the CS and CS-g-PANI substrates. (L) The percentage of the residual mass of the CS and CS-g-PANI hydrogel scaffolds after enzymatic degradation at predetermined biodegradation time points.

### 3.3 Electrical conductivity

Electrical conductivity of swollen hydrogels was obtained using a 2-probe conductivity method shown in Figure 2I. The pure soft and stiff CS hydrogel’s electrical conductivity (as control sample) was 0.19 ±0.041 mS/cm and 0.23± 0.083, respectively. As Figure 2C shows, grafting PANI into CS network significantly improved the electrical conductivity of composite hydrogels. Specifically, the soft and stiff CS-g-PANI samples had conductivities of 7.8 ± 0.35 and 8 ± 0.20 mS/cm, respectively. Note that other studies have also demonstrated that the grafting of polyaniline increased hydrogel electrical conductivity. For example, Zhao et al [46] developed an electroconductive hydrogel based on quaternized chitosan-graft-polyaniline/oxidized dextran and indicated that grafting PANI to hydrogel enhanced the conductivity from 0.43 mS/cm to 1.6 mS/cm. In another investigation, Alizadeh et al showed the ability of oligoaniline for increasing the conductivity of the agarose/alginate/chitosan hydrogel from 0.001 mS/cm to 1 mS/cm [47]. the conductivities of our substrates are at least 5x higher.

### 3.4 Mechanical properties

The shear moduli of CS and CS-g-PANI samples were measured by applying a 2% oscillatory shear strain at 1Hz. For each sample, the shear moduli were collected after the sample had reached a steady state (i.e., after the shear moduli had remained unchanged with time). Figure 2J shows results for the shear moduli averaged over 6 CS samples and 11 CS-g-PANI samples. There were significant differences in the mechanical properties of the soft and stiff CS and CS-g-PANI samples, but no significant differences in stiffness between CS and CS-PANI samples of the same type. The elastic shear moduli of the stiff CS and CS-g-PANI samples at 1 Hz were 40 ± 13 kPa, and 47 ± 16 kPa, respectively, and the viscous shear moduli of those samples were 10.0 ± 6.6 kPa, and 12 ± 4 kPa, respectively. By contrast, the elastic shear moduli of the soft CS and CS-g-PANI samples were 2.8 ± 0.3 kPa and 3.1 ± 0.03 kPa, respectively, and the viscous shear moduli of these samples were 0.24 ± 0.4 kPa, and 0.33 ± 0.3 kPa, respectively. These values of the shear moduli remained almost constant during the time sweep tests, indicating that the samples were mechanically stable.

For both the soft and stiff CS-g-PANI samples, the shear moduli were fairly independent of frequency, at least in the range of 0.01-5 Hz (Supplementary Fig. 2A and 2B). Additionally, the elastic moduli of these samples decreased progressively with increasing strain (Supplementary Fig. 2C and 2D), indicating a shear softening behavior similar to that of soft tissues [48-50]. When the samples were subjected to a 5% steady shear strain, the resulting shear stress gradually decreased with time and saturated at around half of its initial value (Supplementary Fig. 2E and 2F), indicating viscoelastic stress relaxation of the samples. The relaxation appeared to be faster for the stiff CS-g-PANI samples than for the soft ones.

To test if fluid permeation through conducting and non-conducting substrates of the same stiffness might be different, the samples were subjected to a sudden compression, and the resulting axial stress was measured as a function of time after compression. The axial stress relaxation results mainly from the poroelastic effect of solvent being squeezed out of the compressed sample. For the soft CS and CS-g-PANI samples, the axial stress relaxation responses were indistinguishable (Supplementary Fig. 2A), confirming that possible differences in fluid permeation rates cannot account for the differences in cell structure observed on the two substrates. For stiff samples, the CS-g-PANI samples relaxed faster than the CS samples (Supplementary Fig. 2B). However, since the morphologies of cells on the two types of stiff substrates in the absence of electric fields are similar (Figures 4-6), these differences in poroelastic relaxation are unlikely to be relevant to the cells’ response. Since poroelastic rates scale with sample size, the data also show that on the scale of a cell fluid can re-equilibrate through the matrix on a time scale of less than 1 s, confirming that these substrates impose no significant barriers to fluid and therefore soluble metabolite transport.

**Figure 4:**
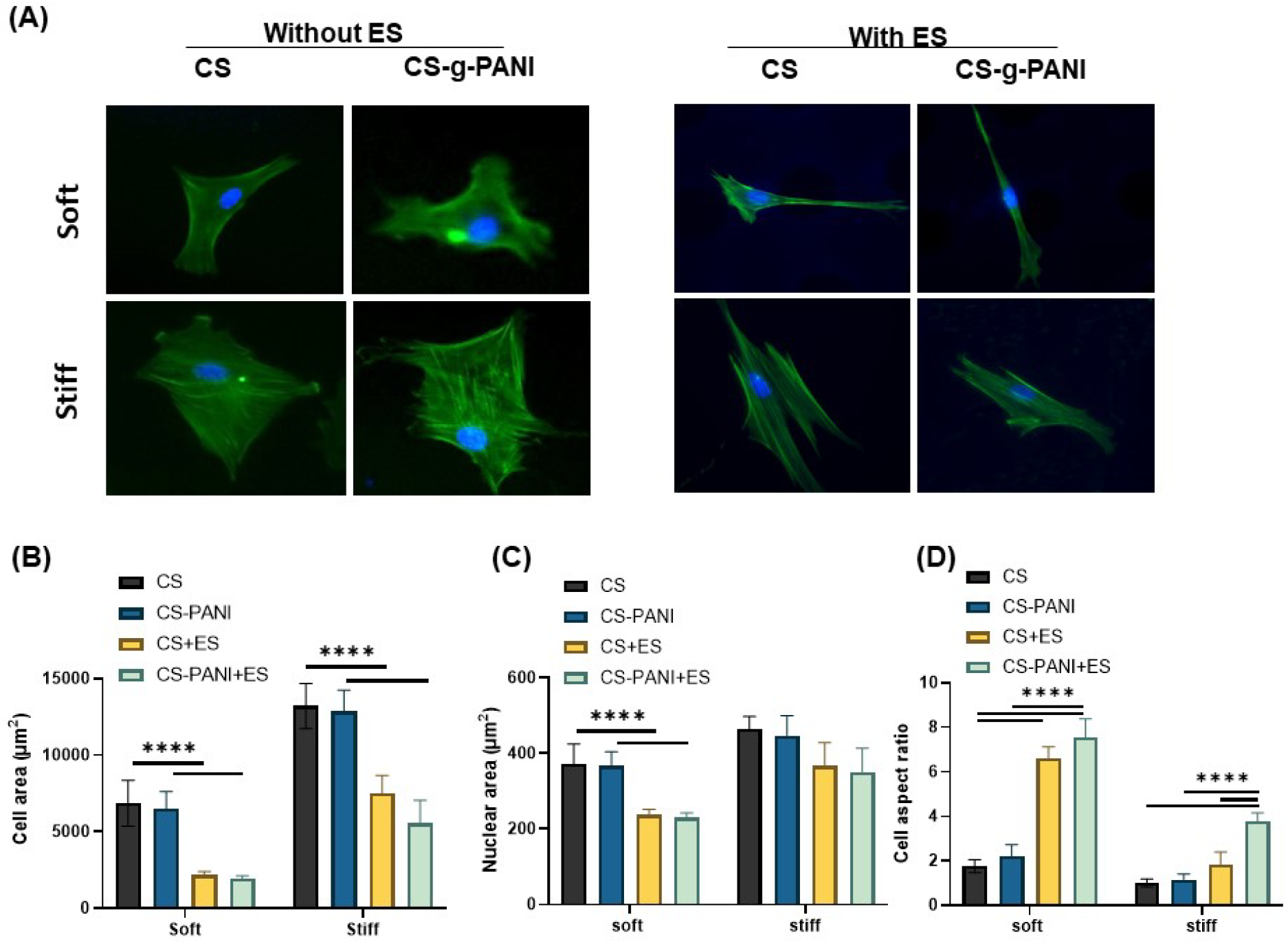
(A) Fluorescence images of hMSCs cultured on CS and CS-g-PANI hydrogel scaffolds. F-actin and nuclei were labeled green and blue, respectively. (Scale bar = 100 μm). (B) Average spread area of cells on each substrate and (C) Cross-sectional area of the cell nuclei on each substrate were quantified using DAPI signals of 150 randomly selected nuclei per independent experiment (n = 3). (D) Cell aspect ratio (ratio of large axis to small axis). Results are shown as mean ± standard deviation. Statistical analysis was performed using two-way ANOVA. Levels of statistical significance (****p < 0.0001) are indicated between days within each substrate.

Supplementary Fig. 4 shows that the electric field does not change the mechanical properties of the conductive substrate. Both the frequency of the free oscillations and the rate of damping, which reflect the shear storage and loss moduli, are indistinguishable before and after application of the 18 V/cm field.

### 3.5 Swelling ratio (SR) and degradation

An equilibrium swelling study was undertaken for hydrogel films after 24 hours of immersion in PBS. Figure 2K displays the equilibrium swelling ratio of CS and CS-g-PANI hydrogels. The SR value for the soft and stiff CS sample was 41% and 23%, respectively, whereas the soft and stiff CS-g-PANI samples had a SR of 27% and 12%, respectively. The decrease in SR for the nanocomposite films is attributed to an increment of hydrophobicity, which hinders water adsorption after grafting PANI chains into the CS network forming a compact and rigid structure. PANI chains not only filled the voids in the CS network but also reduced the pore size of the hydrogels [51]. Therefore, grafting PANI into CS network also reduced the swelling ratio of the conductive CS-g-PANI hydrogels compared to the CS hydrogels.

The degradation rate of scaffolds can directly influence their effects on the tissues, due to the long-term presence of a structure in the body (such a effect can cause a chronic inflammation). Thus, the degradation rate of the hydrogel scaffold should be synchronized with tissue regeneration [52]. The degradation behavior of the prepared hydrogel films was evaluated by analyzing the changes in weight after immersing the samples in collagenase solution in PBS on a rotary shaker (100 rpm). Figure 2L demonstrates biodegradable behaviors of both samples over 4 weeks. In both soft and stiff groups, the weight remaining ratio of the CS-g-PANI substrate was higher compared to CS substrate for a 4-week incubation; therefore, the degradation speed of the CS-g-PANI substrates was lower than that of the CS substrates in each group. After 1 week, the remaining ratios significantly decreased to 71.6±1.8% and 62.4±2% for stiff and soft CS-g-PANI, respectively, and to 64.9 ± 1.2% and 54.2±1.1% for stiff and soft CS, respectively. These ratios then slowly decreased over a 4-week period. This result also agrees with the water swelling ratio data: in the CS-g-PANI hydrogels, water penetration was lower and, thus, the degradation time was longer relative to pure CS hydrogels. These results are similar to the results reported by Ulutürk et al. who showed that there is an inverse relationship between PANI content in CS/PANI hydrogels and the degradation rate of the hydrogels [51].

### 3.6 Cell viability and cell proliferation

The maximal ES intensity that does not have adverse effects on cell viability was first determined. For this purpose, hMSCs were cultured on conductive CS-g-PANI scaffolds with 4 different DC stimuli (50, 100, 150, 200 mV/cm) for a duration of 20 min, and live/dead staining was performed after a further culture for 24 h. The live hMSCs were stained green with calcein acetoxymethyl ester (Calcein-AM), and the dead cells were stained red with ethidium *homodimer*-*1* (EthD-1). Representative images and graphs in Figure 3A show that more than 90% of the hMSCs were alive at 50 and 100 mV/cm, while a reduction of cell viability was observed at 150 and 200 mV/cm. According to these results, 100 mV/cm was considered a proper ES to perform future assessments. Also, after 3, 7, and 14 days of culture in the proliferation medium, cell viability of MSCs on the soft and stiff CS and CS-g-PANI scaffolds was studied for 2 weeks. For this aim, the third passage of primary hMSCs was seeded on the scaffolds at a density of 1 × 10^5^ cells per dish, and live/dead cellular staining was assayed with the assistance of external electrical stimulation (100 mV/cm, 20 min/day). Figure 3B shows a high percentage of hMSCs surviving on both stiff and soft scaffolds under ES, suggesting that the combination of CS and CS-g-PANI with ES (100 mV/cm, 20 min/day) had good biocompatibility after 2 weeks.

**Figure 3:**
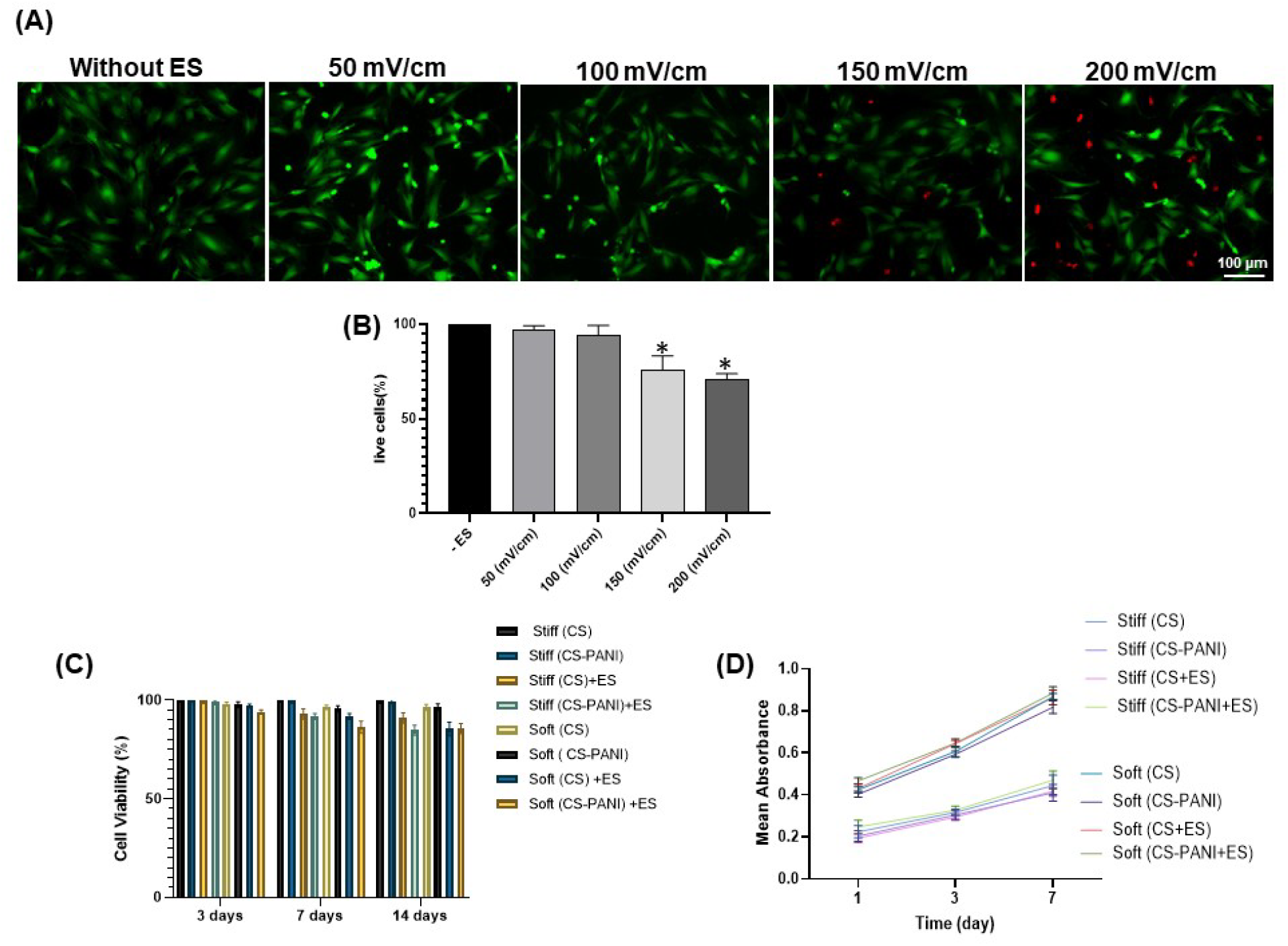
Cytocompatibility of the CS and CS-g-PANI hydrogel scaffolds combined with ES. (A) Live/dead cellular staining of hMSCs seeded on the CS-g-PANI scaffold combined with different ES intensity (50, 100, 150, 200 mV/cm). (B) The graph shows the percent of live cells after each electrical field. * Indicates statistically significant difference (p < 0.05) of electric field treated ES sample w.r.t control without electric field. Live cells are stained green, and dead cells are stained red (scale bar = 100 μm). (C) Cell viability of hMSCs on soft and stiff CS and CS-g-PANI scaffolds with the addition of external electrical stimulation (100 mV/cm) after 3, 7, and 14 days of culture in proliferation medium. (D) Cell proliferation of hMSCs on CS and CS-g-PANI hydrogels scaffolds after 1, 3, and 7 days of culture in the presence of ES. Results are shown as the average values ± standard deviation (n= 3).

The MTT assay was utilized for evaluation of cell proliferation on the various substrates. The proliferation of hMSCs on the scaffolds was determined after cultivation of cells for 1, 3 and 7 days. As illustrated in Figure 3D, the density of hMSCs on both types of scaffolds was higher on stiff samples compared to soft ones at the first day. After 3 and 7 days, for both stiff and soft groups of samples, the proliferation rate of hMSCs cultured on the conductive CS-g-PANI scaffold was slightly lower than that on the pure CS scaffold, but the difference is statistically insignificant. These results showed that incorporation of PANI in the CS hydrogel did not have any adverse effect on cell growth. On the third day, the proliferation rate of hMSCs on the soft CS hydrogels with assistance of ES was higher than on the soft CS hydrogels without ES. The same phenomenon was also observed on both the third and seventh days for both the stiff and soft CS-g-PANI hydrogel groups with/without electrical stimulation. Therefore, applying an electric field on the hydrogels can promote cell proliferation. Jin et al. applied a pulsed electrical potential (100 Hz, 100 mV/ cm) to hMSCs cultured on conductive nanofibers and showed that proliferation of hMSCs cultured with ES was slightly higher compared to that without ES after 7 days of culture [24].

On the other hand, according to the results for each time point of 1, 3 and 7 days, hMSCs grew much faster on stiff samples than soft samples. This observation might be attributed to firmer attachment and the resulting increase in phosphorylated focal adhesion kinase (FAK) caused by higher spreading, as well as reduced apoptosis[53, 54]. Wang et al [53] has shown that hMSCs on stiff gelatin hydroxyphenylpropionic acid (Gtn-HPA) hydrogels (storage modulus = 12800 Pa) exhibited higher proliferation rates compared to soft (Gtn-HPA) hydrogels (storage modulus = 600 Pa).

### 3.7 Cell morphology

In order to determine how substrate stiffness and electrical stimulation affect the attachment and morphological characteristics of hMSCs, actin cytoskeleton labeling with phalloidin was utilized. Fluorescence microscopy was employed to evaluate the cellular spread area, cell aspect ratio, and nuclear area. As Figure 4A shows, on day 1, hMSCs cultured on the stiff CS and CS-PANI substrates displayed the greatest spreading compared to the other samples. The cellular spread area on the soft substrates was approximately half of that on the stiff substrates (Figure 4B). On the other hand, electrically stimulated cells on both the stiff and soft samples had lower spreading area relative to the unstimulated cells cultured on substrates with same stiffness. Therefore, electrically stimulated hMSCs on the soft samples displayed the lowest cell spreading area compared to the other groups. Generally, these data represent the efficacy of electrical stimulation and hydrogel stiffness in controlling cell spreading and contractility.

The cell aspect ratio (defined by the ratio of the major axis to the minor axis) measurement indicated that electrically stimulated hMSCs on soft CS-PANI hydrogels were the most elongated. According to Figure 4C, with assistance of electrical stimulation the hMSCs on the soft CS-g-PANI conductive scaffolds displayed enhanced elongation and oriented growth. In addition, the typical axonal spindle shape of the hMSCs demonstrated that these cells tended to differentiate into neural-like cells after application of the electric field. Rahmani et al. observed spindle-shape morphology of MSCs cultured on a conductive silk fibroin and reduced graphene oxide (SF-rGo) nanofibrous scaffold with 2 electrical impulse models (Current 1:115 V/m, 100-Hz frequency and current 2:115 v/m voltages, 0.1-Hz frequency). In that study, scanning electron microscopy (SEM) images displayed the formation of neuron-like morphology and alignment with electrical impulses [55].

Biophysical signals from the microenvironment are transferred to the nucleus through the cytoskeleton via the LINC complex and can regulate nuclear function [56]. Here, we evaluated the nuclear cross-sectional area as a measure of the stress transferred through cytoskeleton to the nuclei in response to the hydrogel stiffness and ES. The larger tension applied on nuclei including nuclear flattening lead to increased nuclear area. As shown in Figure 4D, hMSCs cultured on stiff hydrogels without ES had the largest nuclear area, which was anticipated considering the greater cellular spreading relative to cells in other substrates. The nuclear area of hMSCs on both the soft and stiff samples indicated a decrease after ES.

### 3.8 Neural Differentiation of hMSCs

Physical cues such as the stiffness, electrical conductivity, and topographic features of substrates direct stem cell behaviors through altering cell shape and rearrangement of cytoskeleton network [57]. It has been shown that substrate stiffness alone can promote a specific lineage differentiation, i.e., MSCs differentiation toward bone, muscle, or neuronal lineage when grown on hard, medium, or soft substrates [58-60]. These results are related to the mechanical properties of the native tissue in which these cells reside. For example, the native brain has 1-4 kPa stiffness, while bone stiffness is at the range of 15-20 MPa [61]. Jih Her et al. reported that hMSCs on type I collagen (Col) and hyaluronic acid (HA) hydrogels at 1 kPa stiffness tended to differentiate into neuronal lineage, while they transformed into glial cells in hydrogels at 10 kPa [62].

On the other hand, the combination of conductive scaffolds with electrical stimulation can influence the cell behavior by changing the membrane potential, altering the ion influx across the cell membrane, or conversion of ligand-receptor formation, which can in turn change intracellular signal transduction pathways [63]. Electrical stimulation can be applied intermittently as well as continuously depending upon the cell type. Zhao et al. demonstrated that a 115 V/m (DC) electric field for 2 h/day can promote directional migration and differentiation of neural precursor cells (NPCs) after culturing for 3 days [64]. It is important to note that hMSCs can form neuron-like cells by activating elements of the neurogenic differentiation pathway [65-68]. However, Ma et al. reported that fully functional neurons cannot be produced by differentiation of MSCs, as they are ectoderm derived [69]. Accordingly, we specifically address immature differentiation by using the term of neuron-like cells in the present context. This process consists of the transformation of multipotent stem cells to committed neuroprogenitor-like cells.

According to above mentioned, in this work, we determined the independent and combined influences of stiffness, conductivity of substrate, and electric field stimulation on the transdifferentiation of hMSCs. For this purpose, hMSCs were precultured on both soft and stiff CS and CS-PANI samples for 24h, followed by electrical stimulation at 100 mV cm^−1^ for 14 days. Figure 5A illustrates the results of immunofluorescence staining for the astrocyte marker (early neural marker) GFAP, the later neural markers MAP2 and NF-H, as well as for the non-neuronal intermediate filament marker vimentin. According to these observations (Figures 5B-E), the glial marker GFAP slightly decreased on the stiff CS and CS-g-PANI scaffolds after electrical stimulation. In contrast, we observed a slight increase of GFAP expression on the soft CS-g-PANI samples. A higher level of MAP2 and NFL expression was seen on the soft conductive CS-g-PANI scaffolds after electrical stimulation. However, the stiff conductive CS-PANI hydrogels in combination with ES exhibited an increment in MAP2 and NFL expression. Also, the expression of both markers was higher on the soft conductive CS-PANI hydrogels relative to the stiff conductive hydrogels. In contrast, the vimentin expression level on hMSCs cultured on the soft and stiff CS and CS-PANI scaffolds significantly decreased. Both with and without ES, the expression of vimentin decreased with decreasing hydrogel stiffness. Neuronal precursors initially express the vimentin (Vim) in vivo, and after the neurons mature, neurofilament proteins replace vimentin [70]. Weng et al. [53] showed that soft gelatinehydroxyphenylpropionic acid (GtneHPA) hydrogels significantly increased MAP2 and NFL expression compared to the same hydrogels with higher stiffness. Substrate stiffness plays a critical role in MSCs differentiation toward neural lineage. Mechanical forces from the cell environment can act through cytoskeleton modification and formation of focal adhesion-stress fiber complex. These formed structures can active cellular signaling pathways, such as RhoA/ROCK signaling, which regulate cell migration, adhesion and differentiation as well as FAK/ERK signaling, MAP kinase signaling that is related to cell proliferation. Balikov et al. demonstrated that both the 0.1 and 0.3 V constant stimulation induced significantly higher expression of β_3_-tubulin and MAP2 in hMSCs seeded on graphene substrate compared to those on the glass control [71]. Electrical fields can also enhance fibroblast growth factor receptor (FGF-R) and its ligand regulated signaling pathway. As a result of electric field application, activation of the FGF signaling pathway can cause increased MAP2 expression [72]. Guo and coworkers reported a significant increase in expression of Tuj1 and GFAP in MSCs cultured on poly(3,4-ethylenedioxythiophene) (PEDOT)− reduced graphene oxide (rGO) hybrid microfibers (with Young’s Modulus › 1 MPa) under electrical stimulation (300 V, 30 μA, 3000 pulses/day for 21 days) [73]. Electrical stimulation is initially sensed by the cell membrane receptors by movement of charges and/or dipoles within the membrane which produces an electric current [74]. Some studies have shown that a DC electric field can polarize a number of cell receptor proteins, including the acetyl choline receptors, VEGF/EGF receptors, and integrins which initiate an intracellular signaling cascade, mediated through src kinase (src), phosphatase and tensin homolog (PTEN), small GTPases, and phosphoinositide kinase pathways [14, 75, 76]. It has been shown by Lim et al. that reduced graphene oxide with a pulsatile electric field can promote the neurogenesis of hMSCs as well as axon and dendrite formation. Focal adhesion kinase (p125fak) has specific interaction with *neural cell adhesion* molecule (NCAM) and the SRC-related tyrosine kinase, which mediate neurite outgrowth [77]. Another hypothesis is that electric field stimulation can change membrane protein conformation, leading eventually to genetic reprogramming. In addition, as cells start to differentiate because of surface mediated genetic reprogramming under electrical stimulation, cells developed a more elongated morphology [78, 79].

**Figure 5:**
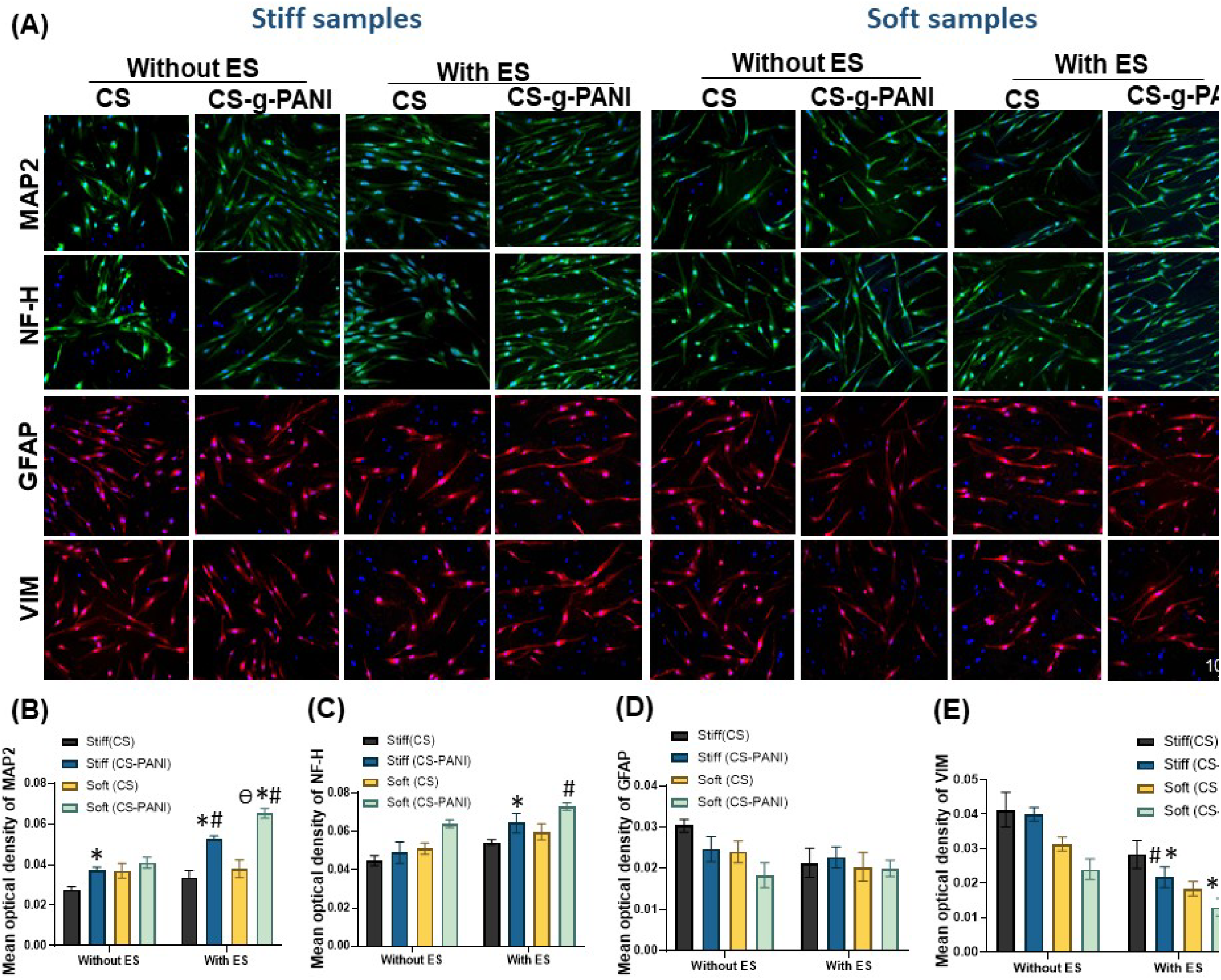
Neural marker expression of neural-like cells differentiated from hMSCs. (A) Fluorescence images of immunostained cells on CS and CS-g-PANI hydrogels with and without electrical stimulation (ES). MAP2 and NF-H are shown in green, GFAP and vimentin are shown in red, and the nuclei are stained with DAPI (blue) (scale bar = 100 μm). (B-E) Statistical analysis of the mean optical density of immunostained images of MAP2, NF-H, GFAP, and vimentin with the assistance of Image J software. * Denotes significance with respect to the corresponding unstimulated counterparts (*p* < 0.05). # Respects statistically significant difference (*p* < 0.05) with respect to electrical stimulation of control cells (on CS sample). θ indicates statistically significant difference (*p* < 0.05) with respect to the corresponding stimulated stiff counterparts.

Remodeling the actin cytoskeleton and producing cell-surface protrusions like filopodia and lamellipodia is a common response of many cells to external electrical fields [80]. To understand the role of ES in cytoskeletal elongation, we also looked at certain measurements of neurite outgrowth on cells with the neuron-like morphology by analyzing fluorescently labeled cells. In the matured neuron MAP2 mostly localizes in neurites. Therefore, the differentiated cells after 14 days were stained for MAP2. Without ES, neurites elongated by 248 ± 42 μm (n =20 cells) in cells on the stiff (CS-g-PANI) substrates, 277± 52 μm in cells on the soft (CS-g-PANI) substrates, 373 ± 30 μm in cells on the stiff (CS-g-PANI) group with ES, and 436±42 μm. in cells on the soft (CS-g-PANI) substrate with ES. Quantitative results are also shown in Figure 6. These results suggest that the pattern of neurite outgrowth in stem cells could be different depending on whether the cells are subjected to ES. The observed morphological change of electrically induced hMSCs on the conductive CS-g-PANI substrates is related to actin cytoskeleton [58] reorganization. Therefore, as the data show, the electrical field and substrate stiffness play an important role in conduction of localization and activation of specific channels or receptors in plasma membrane, [81, 82].

**Figure 6:**
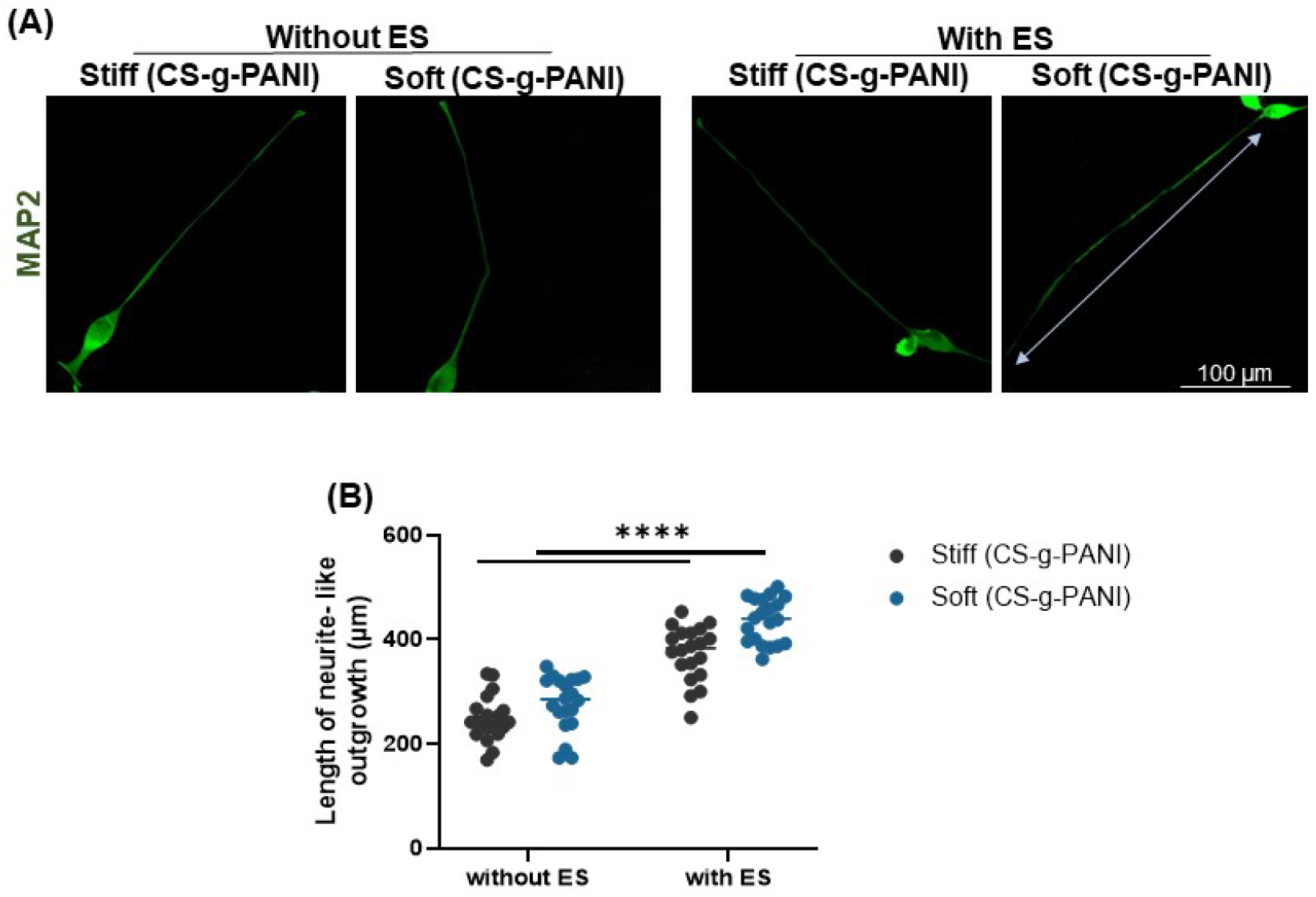
Neurite outgrowth was measured by tracing the total length from the tip to the cell body. All length measurements were performed using the free hand line tool in ImageJ software.

### 3.9 Calcium imaging of differentiated hMSCs

Ca^+2^ signaling has an essential role in neural network development through the regulation of axonal pathfinding, dendritic growth, and specification of neurotransmitter subtype [83], (Figure 7B) [84, 85]. Therefore, we determined the change of intracellular Ca^+2^ dynamics in cells with ES. To determine whether the differentiated hMSCs on the PANI-CS and CS substrates were alive and functionally active, we performed Ca^+2^ imaging. Here, hMSCs were first seeded and differentiated on both the stiff and soft CS-g-PANI substrates for 2 weeks. Afterward, right before ES, differentiated hMSCs on both scaffolds were stained with Fluo-4 AM dye to detect the change of intracellular calcium ion concentration caused by electrical stimuli. Flue 4AM has been used as a membrane-permeable, Ca^+2^ dependent dye, which increases fluorescence on binding of free Ca^+2^ [86]. Previous studies reported that ES can induce a change in Ca^+2^ ion concentration of neuron cells, resulting in the increased fluorescence intensity of Flue 4 AM dye in the neuron [87, 88]. As Figure 7C-D show, after ES, the Fluo-4 fluorescence level of the differentiated hMSCs on both the soft and stiff CS-g-PANI significantly increased compared to cells without electrical stimulation. The left and right fluorescence images in Figures 7C-D display several differentiated cells on the conductive CS-g-PANI samples before and after ES, respectively. The cells marked with the red arrow indicated the enhanced fluorescence intensity after ES, showing the increased cytoplasmic calcium. Figure 8E-F show the relative change Δ *F* / *F* of fluorescence intensity in one cell versus the stimulation time period which shows over ∼ 70% and 85% fluorescence intensity increases in cell on stiff and soft CS-g-PANI substrate by electrical stimuli, respectively. As expected, undifferentiated hMSCs (non-neuronal cells) mostly exhibited nonexcitable behavior and failed to exhibit membrane depolarization after ES treatment. Electrical stimulation through conductive CS-g-PANI scaffolds can activate calcium channels, which leads to Ca^+2^ concentration enrichment. These results imply that CS-g-PANI scaffolds with electrical conductivity properties are a promising strategy to achieve neuronal activation with electrical stimulation.

**Figure 7:**
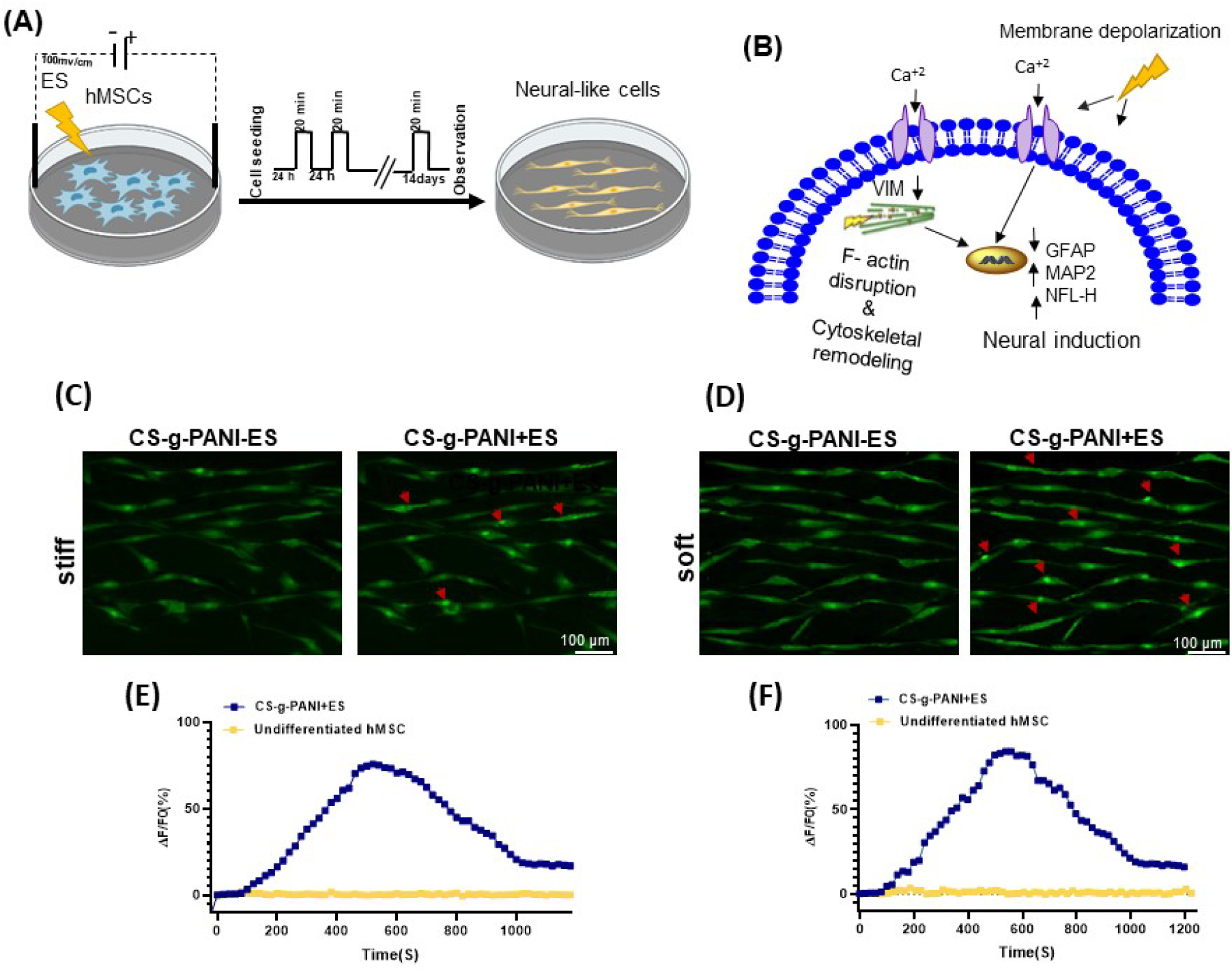
(A) Schematic depicting the morphological alteration and differentiation experienced by hMSCs, cultured on conductive CS-g-PANI substrates and manipulated with ES. Intermittent DC EF stimulation resulted in neural-like morphology. (B) Calcium influx from the extracellular space due to the activation of voltage sensitive calcium channels and Cytoskeletal remodeling because of F-actin disruption are considered to induce neural-like differentiation in electrically stimulated hMSCs. (C) hMSCs on stiff conductive CS-g-PANI substrate were loaded with Fluo-4 AM and observed using fluorescence microscopy. (D) hMSCs on soft conductive CS-g-PANI substrate were loaded with Fluo-4 AM and observed using fluorescence microscopy. An increase in Fluo-4 fluorescence (green) reflects a rise in intracellular Ca^+2^. Red arrow indicates the regions of interest indicating change in Fluo-4AM intensity. (E) Curves referring to the time dependent change in intracellular Ca2+ concentration in undifferentiated (non-stimulated) and stimulated hMSCs on stiff CS-g-PANI substrates. (F) Curves referring to the time dependent change in intracellular Ca2+ concentration in undifferentiated (non-stimulated) and stimulated hMSCs on soft CS-g-PANI substrates. The fluorescence recordings were performed in time-lapse mode at a frequency of one frame in 10 s.

## Conclusion

This study demonstrates that the stiffness of electroconductive scaffolds and electrical stimulation methodologies are both instructive cues for modulation of stem cell commitment towards electrically excitable cells. Specifically, we demonstrated the ability of low-intensity electric field stimulation (100 mV/cm) toward differentiation of hMSCs to neural-like cells seeded on both the soft and stiff conductive CS-g-PANI substrates. Our investigation indicates that CS-g-PANI composite hydrogels exhibit excellent biocompatibility, conductivity, and biodegradability. After electrical stimulation, hMSCs cultured on the soft CS-g-PANI substrates had more elongated morphology with smaller surface area relative to the stiff CS-g-PANI substrates. Moreover, statistically significant upregulation of neural markers in electrically stimulated cells on the soft CS-g-PANI substrates were observed at the protein level. Neuron-like cells on soft CS-g-PANI substrates formed significantly longer and thinner neurites after electrical stimulation. Ca^2+^ imaging results strengthened our finding of differentiation of hMSCs to neural-like cells upon electrical stimulation. Our results suggest the potential applications of soft conductive scaffolds with electric field stimulation to study the differentiation of stromal cells into neural-like cells.

## Supporting Information

**Supplementary Figure 1.**
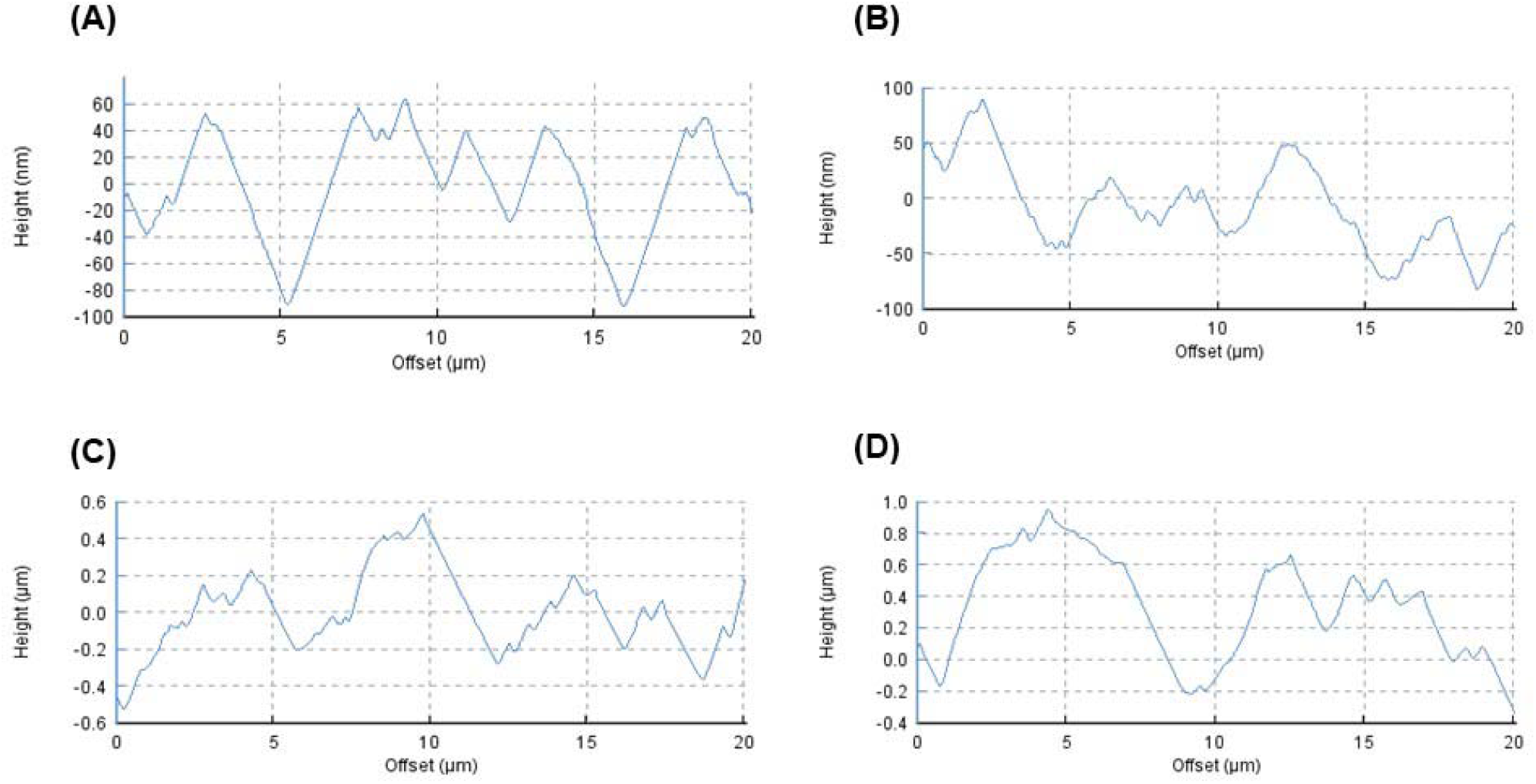
Surface profile of a random horizontal line of stiff CS substrate (A), stiff CS-g-PANI substrate (B), soft CS substrate (C) and soft CS-g-PANI substrate (D).

**Supplementary Figure 2.**
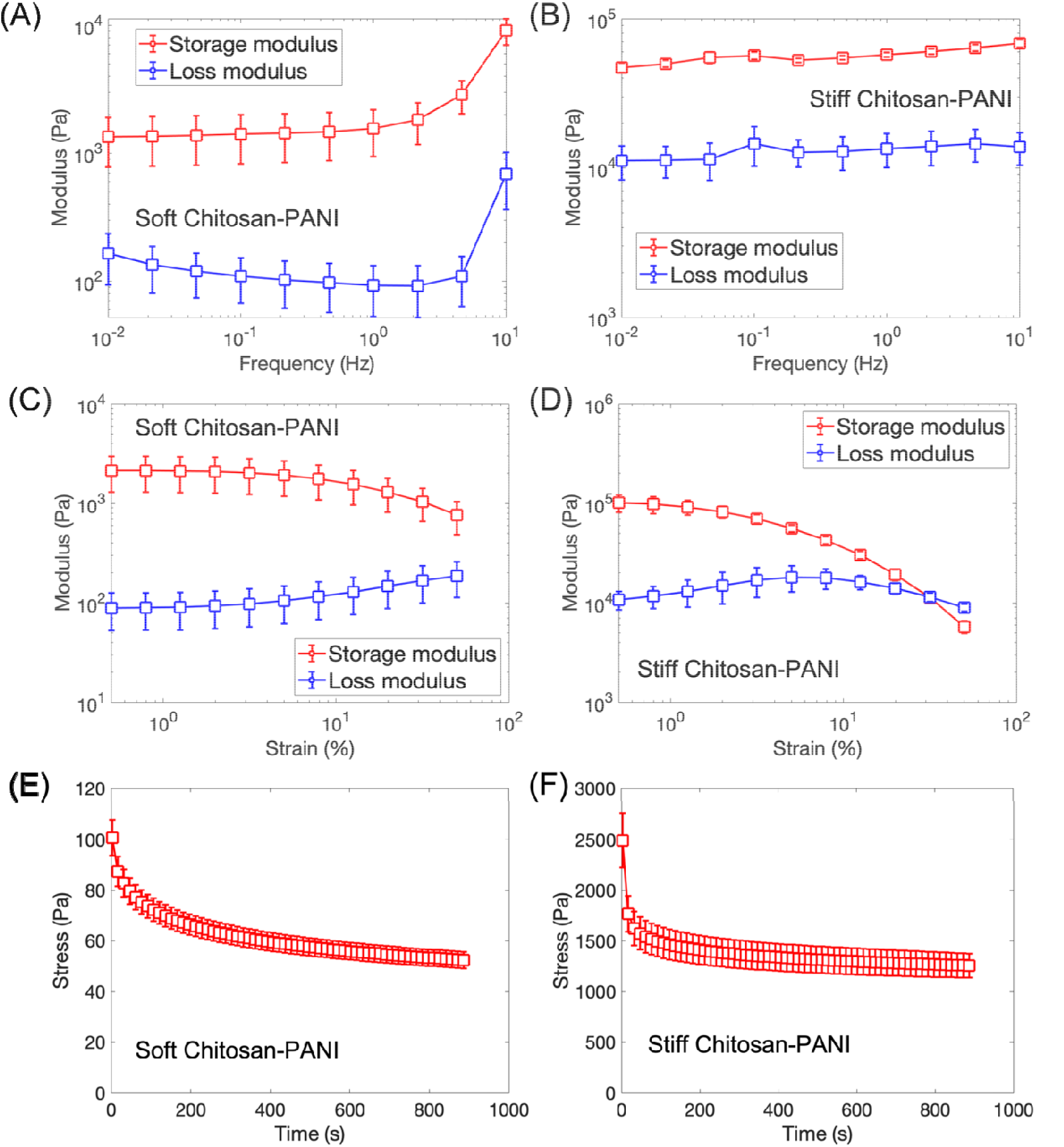
Shear rheology of the CS-g-PANI samples. Results are shown for the frequency sweeps of the (A) soft and (B) stiff samples, the amplitude sweeps of the (C) soft and (D) stiff samples, as well as the shear stress relaxations of the (E) soft and (F) stiff samples.

**Supplementary Figure 3.**
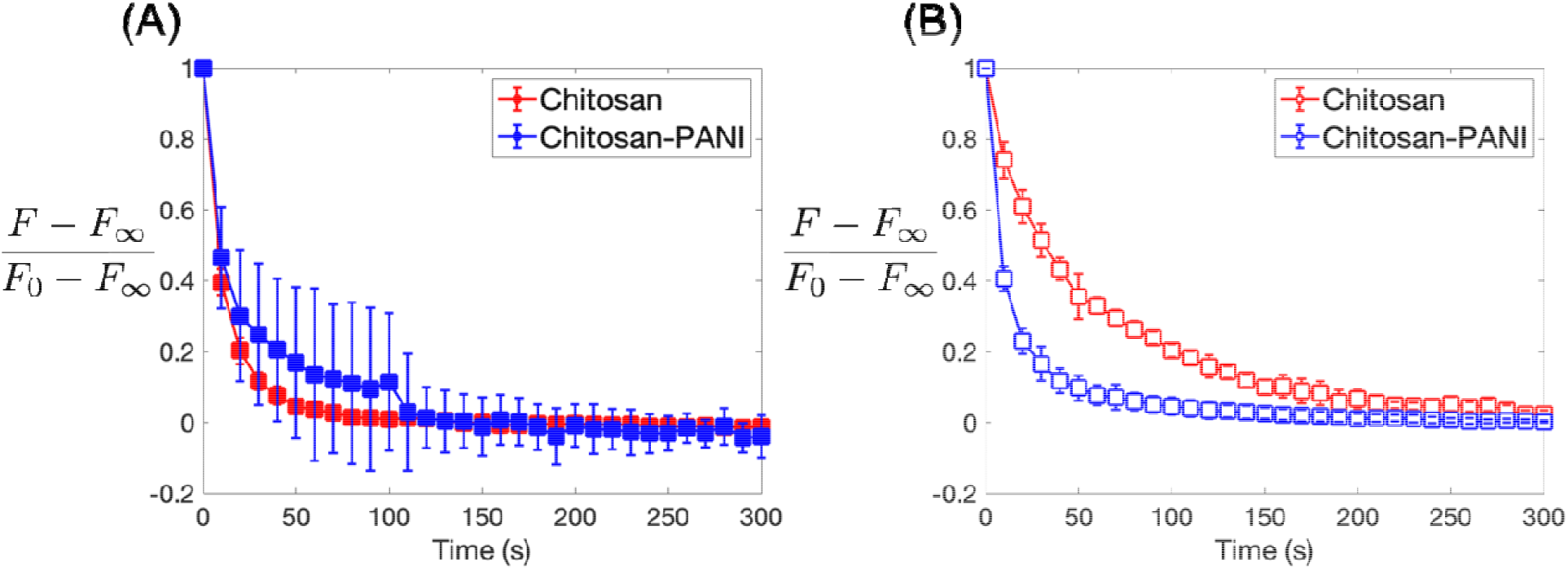
Stress relaxation of the CS and CS-g-PANI substrates in response to compression for (A) soft and (B) stiff samples. The normalized normal force, (, is plotted as a function of time, where is the normal force after the sample had reached equilibrium (s), and is the normal force right after the compression was applied ().

**Supplementary Figure 4.**
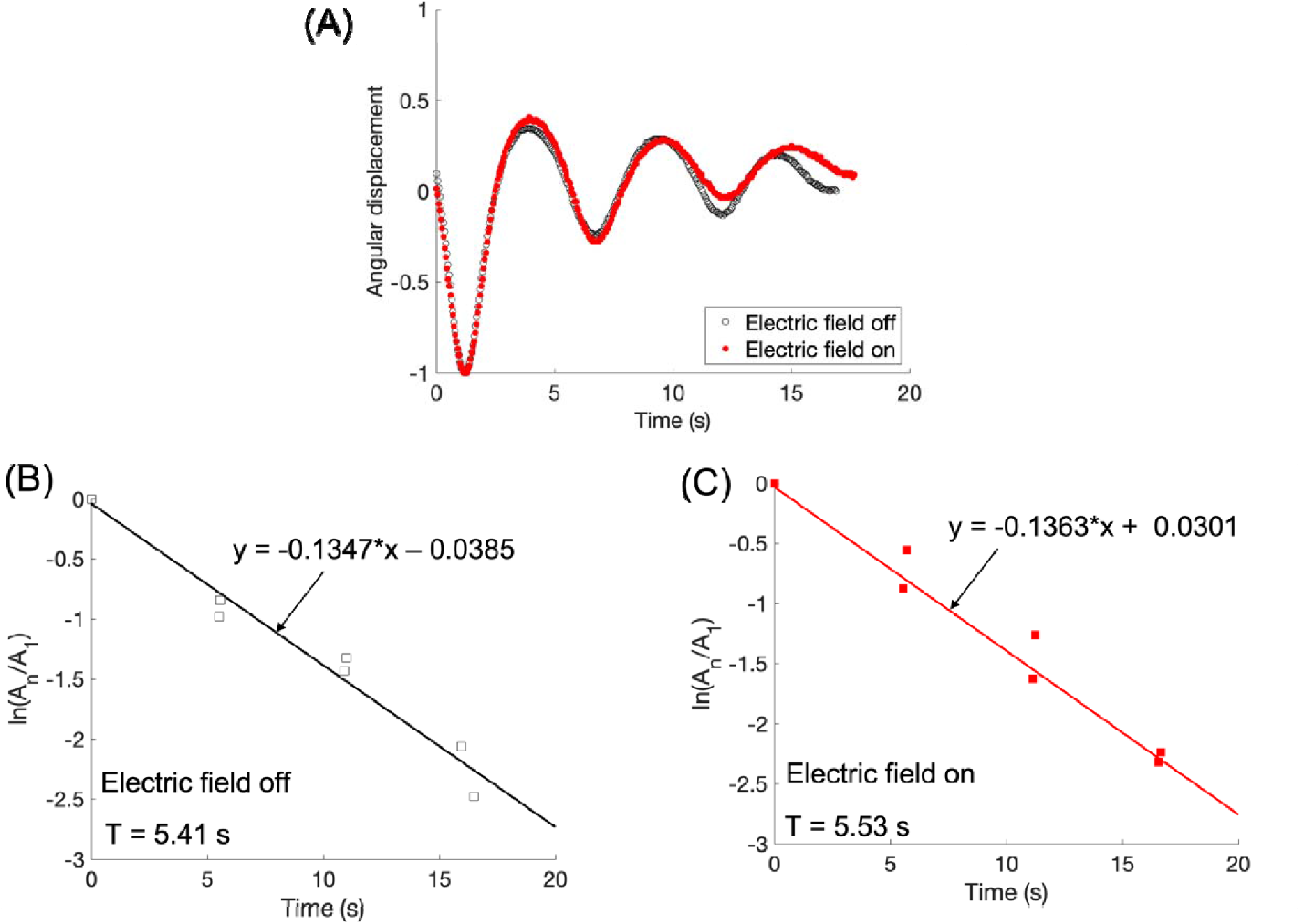
Oscillations of the soft CS-g-PANI samples under a torsion pendulum. (A) Angular displacements of a representative sample during free oscillations before and after applying the electric field. (The results were normalized by the amplitude of the first oscillation). Results are also shown for the logarithmic decay of amplitudes (C) before and (D) after applying the electric field. We found no significant difference in the oscillatory response of the sample before and after applying the electric field. Specifically, the shear storage modulus, G’, and the shear loss modulus, G”, were calculated from the relations in [35] using the torsion pendulum with a moment of inertia of 8693 g cm^2^ and the sample with 8 mm diameter and 2 mm height. The resulting values for G’ are 5850 Pa and 5600 Pa and for G” they are 1050 Pa and 997 Pa, before and after applying the electric field, respectively. These small differences are within the error of the measurement.

## References

1. Dahlin, L.B., et al., Traumatic Peripheral Nerve Injuries: Experimental Models for Repair and Reconstruction, in Animal Models of Neurotrauma. 2019, Springer. p. 169–186.

2. López-Cebral, R., et al., Peripheral nerve injury: current challenges, conventional treatment approaches, and new trends in biomaterials-based regenerative strategies. ACS Biomaterials Science & Engineering, 2017. 3(12): p. 3098–3122.

3. Kornfeld, T., P.M. Vogt, and C.J.W.M.W. Radtke, Nerve grafting for peripheral nerve injuries with extended defect sizes. 2019. 169(9): p. 240–251.

4. Song, B., et al., Application of direct current electric fields to cells and tissues in vitro and modulation of wound electric field in vivo. Nature protocols, 2007. 2(6): p. 1479.

5. Wang, J., et al., In vitro and in vivo studies of electroactive reduced graphene oxide-modified nanofiber scaffolds for peripheral nerve regeneration. Acta biomaterialia, 2019. 84: p. 98–113.

6. Eftekhari, B.S., et al., Surface Topography and Electrical Signaling: Single and Synergistic Effects on Neural Differentiation of Stem Cells. Advanced Functional Materials, 2020: p. 1907792.

7. Margolis, L.B. and S.V. Popov, Generation of cell processes in a high frequency electric field. Bioelectrochemistry and Bioenergetics, 1988. 20(1-3): p. 143–153.

8. Babona-Pilipos, R., et al., Calcium influx differentially regulates migration velocity and directedness in response to electric field application. Experimental cell research, 2018. 368(2): p. 202–214.

9. Balint, R., N.J. Cassidy, and S.H. Cartmell, Electrical stimulation: a novel tool for tissue engineering. Tissue Engineering Part B: Reviews, 2013. 19(1): p. 48–57.

10. Balint, R., N.J. Cassidy, and S.H. Cartmell, Conductive polymers: Towards a smart biomaterial for tissue engineering. Acta biomaterialia, 2014. 10(6): p. 2341–2353.

11. Nair, S.S., S.K. Mishra, and D. Kumar, Recent progress in conductive polymeric materials for biomedical applications. Polymers for Advanced Technologies, 2019. 30(12): p. 2932–2953.

12. Namsheer, K. and C.S.J.R.A. Rout, Conducting polymers: a comprehensive review on recent advances in synthesis, properties and applications. 2021. 11(10): p. 5659–5697.

13. Boni, R., et al., Current and novel polymeric biomaterials for neural tissue engineering. 2018. 25(1): p. 1–21.

14. Zhao, M., et al., Electrical signals control wound healing through phosphatidylinositol-3-OH kinase-γ and PTEN. 2006. 442(7101): p. 457–460.

15. Wrobel, S., et al., In vitro evaluation of cell-seeded chitosan films for peripheral nerve tissue engineering. 2014. 20(17-18): p. 2339–2349.

16. Sadeghi, A., F. Moztarzadeh, and J.A.J.I.j.o.b.m. Mohandesi, Investigating the effect of chitosan on hydrophilicity and bioactivity of conductive electrospun composite scaffold for neural tissue engineering. 2019. 121: p. 625–632.

17. Liu, H., et al., Electrofabrication of flexible and mechanically strong tubular chitosan implants for peripheral nerve regeneration. 2021.

18. Jiang, Z., et al., Rat sciatic nerve regeneration across a 10-mm defect bridged by a chitin/CM-chitosan artificial nerve graft. 2019. 129: p. 997–1005.

19. Islam, M.M., et al., Chitosan based bioactive materials in tissue engineering applications-A review. 2020. 5(1): p. 164–183.

20. Trzaska, K.A. and P. Rameshwar, Dopaminergic neuronal differentiation protocol for human mesenchymal stem cells, in Mesenchymal stem cell assays and applications. 2011, Springer. p. 295-303.

21. Jahan, S., et al., Neurotrophic factor mediated neuronal differentiation of human cord blood mesenchymal stem cells and their applicability to assess the developmental neurotoxicity. Biochemical and biophysical research communications, 2017. 482(4): p. 961–967.

22. Vaca-González, J.J., et al., Effect of electrical stimulation on chondrogenic differentiation of mesenchymal stem cells cultured in hyaluronic acid–Gelatin injectable hydrogels. 2020. 134: p. 107536.

23. Zhang, J., et al., Electrical stimulation of adipose-derived mesenchymal stem cells in conductive scaffolds and the roles of voltage-gated ion channels. 2016. 32: p. 46–56.

24. Jin, L., et al., Synergistic effects of electrical stimulation and aligned nanofibrous microenvironment on growth behavior of mesenchymal stem cells. 2018. 10(22): p. 18543–18550.

25. Eftekhari, B.S., et al., Surface Topography and Electrical Signaling: Single and Synergistic Effects on Neural Differentiation of Stem Cells. 2020. 30(25): p. 1907792.

26. Park, S.J., et al., Neurogenesis is induced by electrical stimulation of human mesenchymal stem cells co-cultured with mature neuronal cells. Macromolecular bioscience, 2015. 15(11): p. 1586–1594.

27. Luo, B., et al., Electrically induced brain-derived neurotrophic factor release from schwann cells. 2014. 92(7): p. 893–903.

28. Mantamadiotis, T., N. Papalexis, and S.J.B. Dworkin, CREB signalling in neural stem/progenitor cells: recent developments and the implications for brain tumour biology. 2012. 34(4): p. 293–300.

29. Eftekhari, B.S., et al., Conductive chitosan/polyaniline hydrogel with cell-imprinted topography as a potential substrate for neural priming of adipose derived stem cells. 2021. 11(26): p. 15795–15807.

30. Engler, A.J., et al., Matrix elasticity directs stem cell lineage specification. Cell, 2006. 126(4): p. 677–89.

31. Kumar, S., et al., Viscoelastic retraction of single living stress fibers and its impact on cell shape, cytoskeletal organization, and extracellular matrix mechanics. 2006. 90(10): p. 3762–3773.

32. Bouzid, T., et al., The LINC complex, mechanotransduction, and mesenchymal stem cell function and fate. 2019. 13(1): p. 1–12.

33. Shivashankar, G.J.C.o.i.c.b., Mechanical regulation of genome architecture and cell-fate decisions. 2019. 56: p. 115–121.

34. Marcasuzaa, P., et al., Chitosan-graft-polyaniline-based hydrogels: elaboration and properties. 2010. 11(6): p. 1684–1691.

35. Janmey, P.A.J.J.o.b. and b. methods, A torsion pendulum for measurement of the viscoelasticity of biopolymers and its application to actin networks. 1991. 22(1): p. 41–53.

36. Plazek, D.J., M. Vrancken, and J.W.J.T.o.t.S.o.R. Berge, A torsion pendulum for dynamic and creep measurements of soft viscoelastic materials. 1958. 2(1): p. 39–51.

37. Abagnale, G., et al., Surface topography guides morphology and spatial patterning of induced pluripotent stem cell colonies. Stem cell reports, 2017. 9(2): p. 654–666.

38. Lv, H., et al., Mechanism of regulation of stem cell differentiation by matrix stiffness. 2015. 6(1): p. 1–11.

39. Yang, K., et al., Electroconductive nanoscale topography for enhanced neuronal differentiation and electrophysiological maturation of human neural stem cells. Nanoscale, 2017. 9(47): p. 18737–18752.

40. Jin, G., K.J.M.S. Li, and E. C, The electrically conductive scaffold as the skeleton of stem cell niche in regenerative medicine. 2014. 45: p. 671–681.

41. Teixeira, A.I., et al., The promotion of neuronal maturation on soft substrates. 2009. 30(27): p. 4567–4572.

42. Sthanam, L.K., et al., Initial priming on soft substrates enhances subsequent topography-induced neuronal differentiation in ESCs but not in MSCs. 2018. 5(1): p. 180–192.

43. Yavuz, A.G., A. Uygun, and V.R.J.C.P. Bhethanabotla, Substituted polyaniline/chitosan composites: Synthesis and characterization. 2009. 75(3): p. 448–453.

44. Karthik, R. and S.J.I.j.o.b.m. Meenakshi, Facile synthesis of cross linked-chitosan–grafted-polyaniline composite and its Cr (VI) uptake studies. 2014. 67: p. 210–219.

45. Sahnoun, S. and M.J.I.j.o.b.m. Boutahala, Adsorption removal of tartrazine by chitosan/polyaniline composite: kinetics and equilibrium studies. 2018. 114: p. 1345–1353.

46. Zhao, X., et al., Antibacterial and conductive injectable hydrogels based on quaternized chitosan-graft-polyaniline/oxidized dextran for tissue engineering. 2015. 26: p. 236–248.

47. Alizadeh, R., et al., Conductive hydrogels based on agarose/alginate/chitosan for neural disorder therapy. 2019. 224: p. 115161.

48. van Oosten, A.S., et al., Emergence of tissue-like mechanics from fibrous networks confined by close-packed cells. 2019. 573(7772): p. 96–101.

49. Mihai, L.A., et al., A comparison of hyperelastic constitutive models applicable to brain and fat tissues. 2015. 12(110): p. 20150486.

50. Song, D., et al., Cell-induced confinement effects in soft tissue mechanics. 2021. 129(14): p. 140901.

51. Ulutürk, C. and N.J.C.p. Alemdar, Electroconductive 3D polymeric network production by using polyaniline/chitosan-based hydrogel. 2018. 193: p. 307–315.

52. Zarrintaj, P., et al., A facile route to the synthesis of anilinic electroactive colloidal hydrogels for neural tissue engineering applications. 2018. 516: p. 57–66.

53. Wang, L.-S., et al., The role of stiffness of gelatin–hydroxyphenylpropionic acid hydrogels formed by enzyme-mediated crosslinking on the differentiation of human mesenchymal stem cell. 2010. 31(33): p. 8608–8616.

54. Hsiong, S.X., et al., Differentiation stage alters matrix control of stem cells. 2008. 85(1): p. 145–156.

55. Rahmani, A., et al., Conductive electrospun scaffolds with electrical stimulation for neural differentiation of conjunctiva mesenchymal stem cells. 2019. 43(8): p. 780–790.

56. Kalukula, Y., et al., Mechanics and functional consequences of nuclear deformations. 2022: p. 1–20.

57. Wang, S., et al., The effect of physical cues of biomaterial scaffolds on stem cell behavior. 2021. 10(3): p. 2001244.

58. Engler, A.J., et al., Matrix elasticity directs stem cell lineage specification. 2006. 126(4): p. 677–689.

59. Higuchi, A., et al., Physical cues of biomaterials guide stem cell differentiation fate. 2013. 113(5): p. 3297–3328.

60. Uz, M., et al., Advances in controlling differentiation of adult stem cells for peripheral nerve regeneration. 2018. 7(14): p. 1701046.

61. Dey, K., et al., Progress in the mechanical modulation of cell functions in tissue engineering. 2020. 8(24): p. 7033–7081.

62. Her, G.J., et al., Control of three-dimensional substrate stiffness to manipulate mesenchymal stem cell fate toward neuronal or glial lineages. 2013. 9(2): p. 5170–5180.

63. Liu, Z., et al., Electroactive Biomaterials and Systems for Cell Fate Determination and Tissue Regeneration: Design and Applications. 2021: p. 2007429.

64. Zhao, H., et al., Specific intensity direct current (DC) electric field improves neural stem cell migration and enhances differentiation towards βIII-tubulin+ neurons. 2015. 10(6): p. e0129625.

65. Tsai, H.-L., et al., Wnts enhance neurotrophin-induced neuronal differentiation in adult bone-marrow-derived mesenchymal stem cells via canonical and noncanonical signaling pathways. 2014. 9(8): p. e104937.

66. Thrivikraman, G., G. Madras, and B.J.B. Basu, Electrically driven intracellular and extracellular nanomanipulators evoke neurogenic/cardiomyogenic differentiation in human mesenchymal stem cells. 2016. 77: p. 26–43.

67. Li, H., et al., Micropatterning extracellular matrix proteins on electrospun fibrous substrate promote human mesenchymal stem cell differentiation toward neurogenic lineage. 2016. 8(1): p. 563–573.

68. Sallam, A., et al., In vitro differentiation of human bone marrow stromal cells into neural precursor cells using small molecules. 2021. 363: p. 109340.

69. Ma, K., et al., Generation of neural stem cell-like cells from bone marrow-derived human mesenchymal stem cells. 2011. 33(10): p. 1083–1093.

70. Yabe, J.T., et al., Regulation of the transition from vimentin to neurofilaments during neuronal differentiation. 2003. 56(3): p. 193–205.

71. Balikov, D.A., et al., Directing lineage specification of human mesenchymal stem cells by decoupling electrical stimulation and physical patterning on unmodified graphene. 2016. 8(28): p. 13730–13739.

72. Titushkin, I. and M.J.B.j. Cho, Regulation of cell cytoskeleton and membrane mechanics by electric field: role of linker proteins. 2009. 96(2): p. 717–728.

73. Guo, W., et al., Self-powered electrical stimulation for enhancing neural differentiation of mesenchymal stem cells on graphene–poly (3, 4-ethylenedioxythiophene) hybrid microfibers. 2016. 10(5): p. 5086–5095.

74. Pullar, C.E., et al., β4 integrin and epidermal growth factor coordinately regulate electric field-mediated directional migration via Rac1. 2006. 17(11): p. 4925–4935.

75. Tsai, C.H., B.J. Lin, and P.H.G.J.J.o.O.R. Chao, α2β1 integrin and RhoA mediates electric field-induced ligament fibroblast migration directionality. 2013. 31(2): p. 322–327.

76. Lin, B.-j., et al., Lipid rafts sense and direct electric field-induced migration. 2017. 114(32): p. 8568–8573.

77. Lim, K.T., et al., Pulsed-electromagnetic-field-assisted reduced graphene oxide substrates for multidifferentiation of human mesenchymal stem cells. 2016. 5(16): p. 2069–2079.

78. Aaron, R.K., et al., Stimulation of growth factor synthesis by electric and electromagnetic fields. 2004. 419: p. 30–37.

79. Pullar, C.E., The physiology of bioelectricity in development, tissue regeneration and cancer. 2016: CRC press.

80. Lee, S.-L., et al., Physically-induced cytoskeleton remodeling of cells in three-dimensional culture. 2012. 7(12): p. e45512.

81. Wu, Y., et al., Conductive micropatterned polyurethane films as tissue engineering scaffolds for Schwann cells and PC12 cells. 2018. 518: p. 252–262.

82. Ke, Y., et al., Smart windows: electro-, thermo-, mechano-, photochromics, and beyond. 2019. 9(39): p. 1902066.

83. Rosenberg, S.S. and N.C.J.C.S.H.p.i.b. Spitzer, Calcium signaling in neuronal development. 2011. 3(10): p. a004259.

84. Tonelli, F.M., et al., Stem cells and calcium signaling. 2012: p. 891–916.

85. Hao, B., et al., The role of Ca2+ signaling on the self-renewal and neural differentiation of embryonic stem cells (ESCs). 2016. 59(2-3): p. 67–74.

86. Gee, K.R., et al., Chemical and physiological characterization of fluo-4 Ca2+-indicator dyes. 2000. 27(2): p. 97–106.

87. Chou, N., et al., A Multimodal Multi-Shank Fluorescence Neural Probe for Cell-Type-Specific Electrophysiology in Multiple Regions across a Neural Circuit. 2021: p. 2103564.

88. Wu, Y., et al., Photoconductive micro/nanoscale interfaces of a semiconducting polymer for wireless stimulation of neuron-like cells. 2019. 11(5): p. 4833–4841.

